# On-person adaptive evolution of *Staphylococcus aureus* during treatment for atopic dermatitis

**DOI:** 10.1101/2021.03.24.436824

**Authors:** Felix M. Key, Veda D. Khadka, Carolina Romo-González, Kimbria J. Blake, Liwen Deng, Tucker C. Lynn, Jean C. Lee, Isaac M. Chiu, Maria Teresa García-Romero, Tami D. Lieberman

**Affiliations:** Institute for Medical Engineering and Science, Massachusetts Institute of Technology, Cambridge, MA, USA; Department of Civil and Environmental Engineering, Massachusetts Institute of Technology, Cambridge, MA, USA; Experimental Bacteriology Laboratory, National Institute for Pediatrics, Mexico City, Mexico; Department of Immunology, Blavatnik Institute, Harvard Medical School, Boston, MA, USA; Division of Infectious Disease, Department of Medicine, Brigham and Women’s Hospital and Harvard Medical School, Boston, MA, USA; Department of Dermatology, National Institute for Pediatrics, Mexico City, Mexico; Broad Institute, Massachusetts Institute of Technology, Cambridge, MA, USA; Ragon Institute, Massachusetts Institute of Technology, Cambridge, MA, USA

## Abstract

Genetic variation among bacterial strains can contribute to heterogeneity in the severity of chronic inflammatory diseases ^1,2^, but the degree of variation created by *de novo* mutation during colonization is not well understood. The inflamed skin of people with atopic dermatitis (AD) is frequently colonized with *Staphylococcus aureus*, an opportunistic pathogen associated with both asymptomatic colonization of nasal passages and invasive disease ^3–6^. While genetic risk and barrier disruption are critical to AD initiation ^7,8^, *S. aureus* colonization is thought to worsen disease severity by promoting skin damage^9 1,4,5,10^. Here we show, from tracking 23 children treated for AD over 9 months, that *S. aureus* adapts via *de novo* mutations during colonization. Patients’ *S. aureus* populations are typically dominated by a single lineage, with infrequent invasion by distant lineages. Variants emerge within each lineage with mutation accumulation rates similar to *S. aureus* in other contexts. Some of these variants replace their ancestors across the body within months, with signatures of adaptive, rather than neutral, forces. Most strikingly, the capsule synthesis gene *capD* obtained four parallel mutations within one patient and was involved in mutational sweeps in multiple patients. We confirm that selection for *capD* negativity is common in AD, but not in other contexts, via reanalysis of public *S. aureus* genomes from 276 people. Our finding of disease-specific selection raises the possibility that adaptation of pathobionts during colonization prolongs the positive feedback cycle of inflammation.

## Main

During colonization of human microbiomes, bacteria acquire adaptive mutations that enhance their ability to survive in the human environment, resist antibiotics, and outcompete other strains ^11–14^. While these *de novo* mutations rise in frequency due to the survival advantage they provide to the bacteria, their emergence may impact host metabolism, immune homeostasis, or microbiome dynamics. Understanding the tempo and consequences of variations across bacterial genomes is of particular importance for complex inflammatory diseases like atopic dermatitis and inflammatory bowel diseases, for which the causative role of the microbiome has been hard to pin down ^15,16^. While recent studies have identified bacterial strains associated with inflammatory states ^1,17,18^, classic metagenomic approaches do not provide the resolution to robustly identify individual mutations emerging in disease states. As a result, the potential impact of *de novo* microbiome mutations on complex diseases is poorly understood.

Atopic dermatitis (AD) is one such chronic inflammatory skin disease with strong microbial associations and a complex etiology. AD transiently affects up to 20% of people during their lifetime ^19^ and is particularly prominent among children, who develop itchy patches of inflamed skin, typically located on the cubital and popliteal fossae (inside of elbows and backs of knees)^20^. Genetic and environmental defects in barrier function have been associated with AD, but are insufficient to explain the variation in disease development and response to treatment ^21^. Notably, symptomatic AD skin of children and adults is usually colonized by the opportunistic pathogen *Staphylococcus aureus*, with abundance proportional to disease severity ^4–6,9,22,23^; this species is usually lowly abundant on healthy skin ^24,25^. Its native reservoir is thought to be the nares, where it asymptomatically colonizes 30% of healthy individuals ^24^. However, *S. aureus* also causes a variety of human infections of the skin, bloodstream, lung, and bone ^3^.

*S. aureus* sequence types vary in virulence potential, in the antibiotic resistance cassettes they carry, and the disease contexts in which they are most often found (e.g. hospital or community associated)^3,26^, yet no sequence types have been robustly associated with AD ^10^. A recent study has suggested that *de novo* mutations in a key quorum sensing pathway reduce the risk for healthy babies to develop AD ^27^--supporting a possible role for *S. aureus* mutations beyond the strain level in AD. While *de novo* mutations occurring in *S. aureus* in young children with AD have been observed ^28^, the fate of these mutations over time and their consequences have not been characterized.

### Longitudinal tracking of *S. aureus* evolution in AD

Here, we use longitudinal sampling and culture-based whole-genome sequencing to identify mutations acquired by *S. aureus* that emerge under natural selection on individual people. We conducted a prospective, longitudinal study of 23 children (age 5 to 15 years old) in Mexico with moderate to severe AD at 5 visits over the course of approximately 9 months (**Figure 1a**). Patients were treated for AD with the standard of care, including topical steroids, emollients, and some with bleach baths. Treatment was modified at each visit according to severity, though patients were not closely following before the final visit, typically about 6 months (**Supplementary Table 1**). Patients were asked not to take antibiotics during the initial 3 month period, but chart review identified that 3 of the 23 patients were administered antibiotics in this initial period due to medical need, and 5 took antibiotics after this initial period; these patients were included in the analysis, and antibiotic usage is noted (see **Supplementary Table 1** for details). During each visit, AD severity was assessed using the SCORing Atopic Dermatitis (SCORAD) scale ^29^, and swabs were collected from seven affected and unaffected skin sites, including cubital and popliteal fossae, forearms and the nares (**Figure 1a**). From each of 225 swabs that yielded growth resembling *S. aureus*, we picked up to 10 single-colony isolates for subculture and sequencing (Methods), resulting in 1,499 *S. aureus* whole genomes.

**Figure 1:**
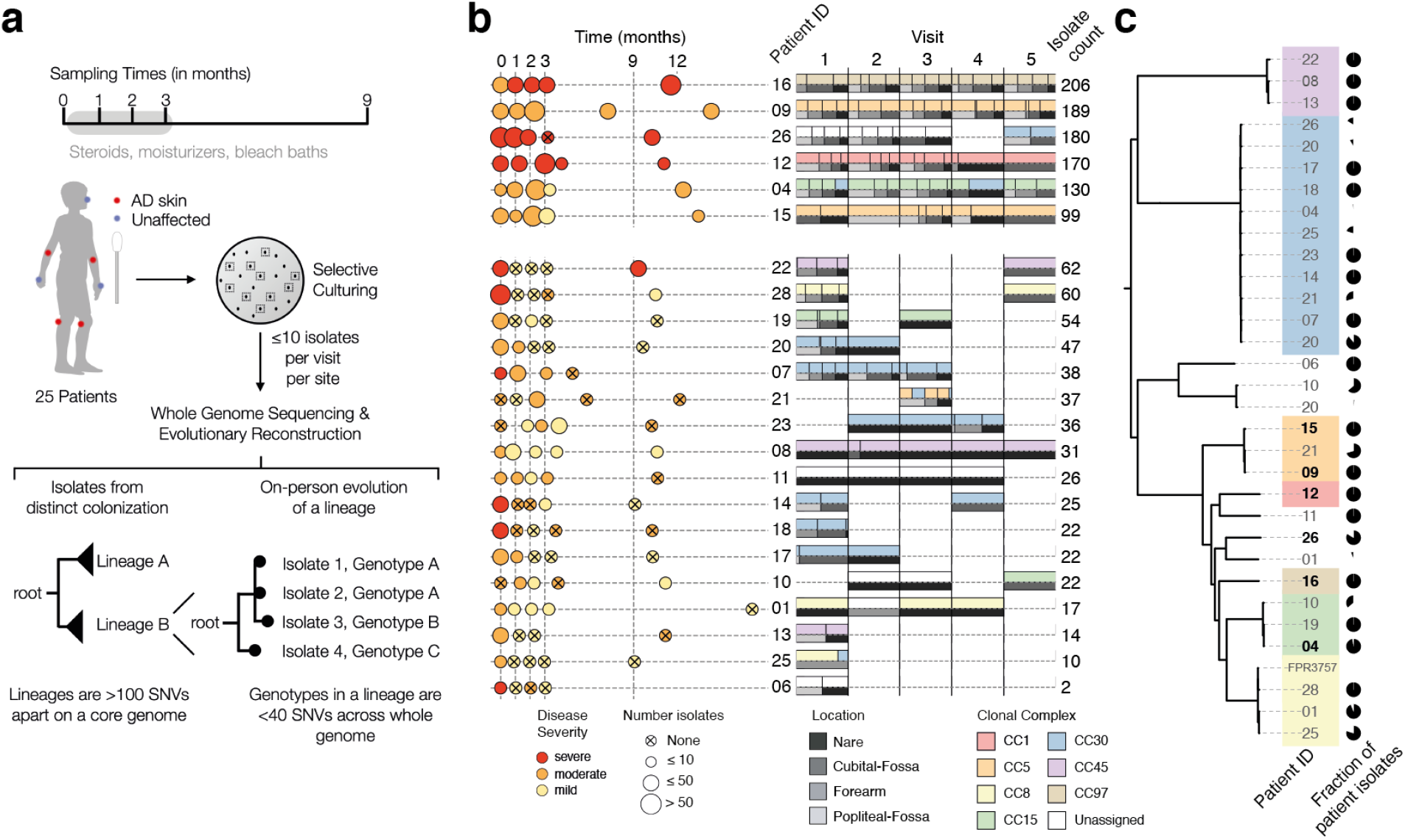
The *S. aureus* population on each child with atopic dermatitis (AD) is dominated by a single, stable, patient-specific lineage. **a,** Swab samples were collected from each of 7 sites during each of 5 visits, including from nares and areas rarely affected by AD in children (forearms). Patient treatment for AD with standard treatment of care, with or without bleach baths, was monitored closely between the first four visits (Methods). From each swab, up to 10 colonies were cultured and processed for whole-genome sequencing. We then grouped isolates from each patient into lineages such that isolates from the same lineage are fewer than 100 mutations across the core genome, and performed more detailed evolutionary analysis within each lineage. **b,** Left panel: Sampling time, mean disease severity at visit based on SCORAD (mild < 25 ≤ moderate < 50 ≤ severe), and number of isolates per visit (dot size) are shown for each patient. Some sampling visits deviated from the anticipated schedule (vertical dashed line), spacing shown reflects the actual sampling schedule achieved (Methods). Right panel: Pairwise stacked bar plots show the inferred clonal complex (top) and sampling location (bottom) for all isolates at each visit. Six patients with ≥ 99 isolates were analyzed with the most detail. **c,** Phylogenetic tree showing the relationship between all 1,499 *S. aureus* isolates sequenced, generated using 60,973 SNVs from a reference-based approach (Methods) and labeled by patient of origin. Pie charts indicate the fraction of patient isolates that come from each lineage, showing most patients have a single major lineage. Patient 20 was colonized by three lineages, two of which are part of the same clonal complex (CC30). Lineages are colored according to their assigned clonal complex (Methods), highlighting that strains colonizing AD patients come from a wide variety of global diversity.

To understand the minimal number of independent colonizations of *S. aureus* onto each child, we first clustered each patient’s isolates into lineages, where each lineage is composed of isolates separated by < 100 mutations across the core genome (**Extended Data Figure 1**). Given the known rates of mutation accumulation in *S. aureus* (~8 mutations/whole genome/yr) ^30–35^, isolates from distinct lineages are too distant to have arisen from a single colonization and subsequent on-person diversification.

Most patients were stably colonized by a single major lineage, though we recovered minority lineages from 7 patients and major lineage replacements in 2 of 19 patients with *S. aureus* recovered at multiple timepoints (**Figure 1b; Supplementary Table 2**). Overall, detected lineages span the diversity of the *S. aureus* species (**Figure 1c**)^36^. The largest fraction of lineages are part of clonal complex 30 (33% of lineages; 15% of isolates), in contrast with a previous finding of clonal complex 1 dominance among people with AD in the UK ^28^.

The number of isolates recovered and from which sufficient sequencing reads were obtained is variable across visits and correlates with both disease severity and *S. aureus* relative abundance inferred from 16S rRNA sequencing ^23^ (0-68 isolates/visit; r^2^=0.36 and r^2^=0.16, respectively; **Figure 1b** and **Extended Data Figure 2**). Six patients provided large numbers of isolates amenable to more in-depth quantitative analyses (99-206 isolates/patient; **Figure 1b**).

### *S. aureus* mutants sweep across the body

To analyze the causes and consequences of bacterial mutation on each person, we next focused on the genetic variation within each stably colonizing lineage. For each patient’s major lineage, we generated a lineage-specific *S. aureus* assembly, used a rigorous alignment-based approach to identify single-nucleotide variants (SNVs), and built phylogenetic trees that illustrated mutation accumulation on the patient (**Figure 2a,** Methods). In many cases, multiple colonies from a timepoint were indistinguishable across their entire genome; we refer to each group of isolates with identical genomes as a ‘genotype’.

**Figure 2.**
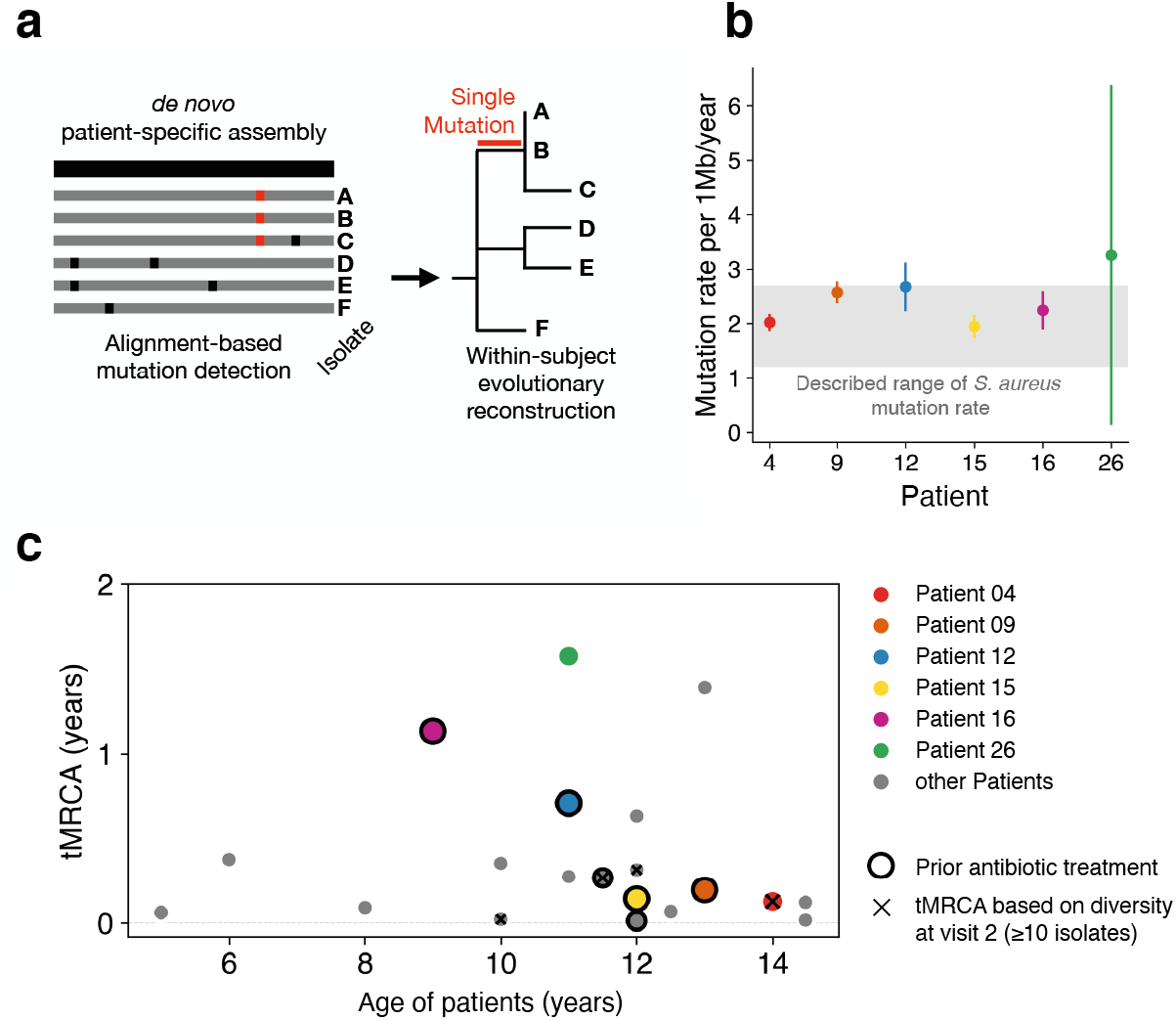
Despite high rates of mutation accumulation on each person, the intralineage diversity on each person is limited. **a,** We built a *de novo* reference genome for each patient (black) and called *de novo* mutations by aligning raw reads from each sample individually to this reference genome, allowing the building of fine-resolution phylogenetic trees (Methods). **b,** The median molecular clock rate (Linear least-squares regression) is shown for the six patients with the most isolates. Error bars indicate 95% confidence intervals. The gray region indicates rates previously reported for *S. aureus* in other contexts ^30–35^. **c,** For each patient with ≥10 isolates at a visit (20 patients), the inferred time elapsed since the population’s most recent common ancestor (tMRCA; Methods) is plotted relative to the patient’s age. Low values of tMRCA indicate a recent single-cell bottleneck, and are found across patients regardless of antibiotic usage within the past 6 months (black outline) or age. The earliest such visit from each patient was used; x’s indicate patients for which visit 2 was used. Mutational distances were converted to time using the calculated molecular clocks for the 6 highlighted patients; for all others, the median rate was used.

In theory, the inflammatory environment on AD skin might induce an elevated mutation rate. We estimated the molecular clock of the lineages colonizing the six patients with the most isolates. For five of these patients’ lineages, molecular clock estimates were consistent with those of *S. aureus* in other contexts, including healthy carriage and in invasive infections (1.2 - 2.7 × 10^-6^ substitutions/site/year ^30–35^; **Figure 2b**). Patient 26’s major lineage had a substantially higher accumulation mutation rate of 3.3 × 10^-6^ substitutions/site/year (CI95 = 0.1 - 6.4 × 10^-6^ substitutions/site/year), though this difference was not significant. While this non-significantly elevated mutation accumulation rate could plausibly reflect environmental conditions that induced higher mutagenesis or a defect in DNA repair not apparent in our genomic analyses, we do not observe any significant differences in mutational spectra across patients (**Extended Data Figure 3**). Taken as a whole, these results suggest that AD-associated inflammation does not greatly increase *S. aureus* mutation rates.

Accumulation of mutations over time can produce two different patterns of on-person evolution: diversification into long co-existing genotypes or within-lineage genotypic replacement ^37,38^. To determine which of these patterns was more common, we calculated the time since each major lineage’s isolates shared a single-celled ancestor (time to the most recent common ancestor, tMRCA). We found that all 16 patients with sufficient data at the first visit (≥10 isolates) had low values of tMRCA indicating seeding of the population from a single genotype within the past few years (tMRCA <1.6 years; **Figure 2c**). We did not find a dependence on recent antibiotic usage. In addition, similarly low values of tMRCA were found for samples collected after initiation of study-related treatment (major lineages where visit 1 yielded ≤10 isolates; **Figure 2c**) and in a reanalysis of 8 AD patients from an unrelated study from which ≥10 isolates were collected at a single timepoint (**Extended Data Figure 4**). Together with our observations of lineage-level stability (**Figure 1b**), these analyses suggests that within-lineage genotypic replacement is a universal aspect of *S. aureus* growing on treated AD skin.

To better understand within-lineage replacement, we examined within-patient phylogenies for all patients (**Figure 3; Extended Data Figure 5–6)**. Visualizing the temporal evolutionary dynamics using Muller diagrams revealed a common pattern in which genotypes with newly acquired mutations repeatedly replaced existing diversity (**Figure 3a-f**). These replacements spanned the entire body; the spread of new mutations was not spatially restricted and included the natural habitat of the nose (**Figure 3g-h; Extended Data Figure 5–6**). While some of these sweeps overlapped with antibiotic administration between samplings, many intervals without antibiotic usage demonstrate similarly dramatic changes in mutational composition, further supporting the presence an antibiotic-independent mechanism of genotypic sweeps.

**Figure 3.**
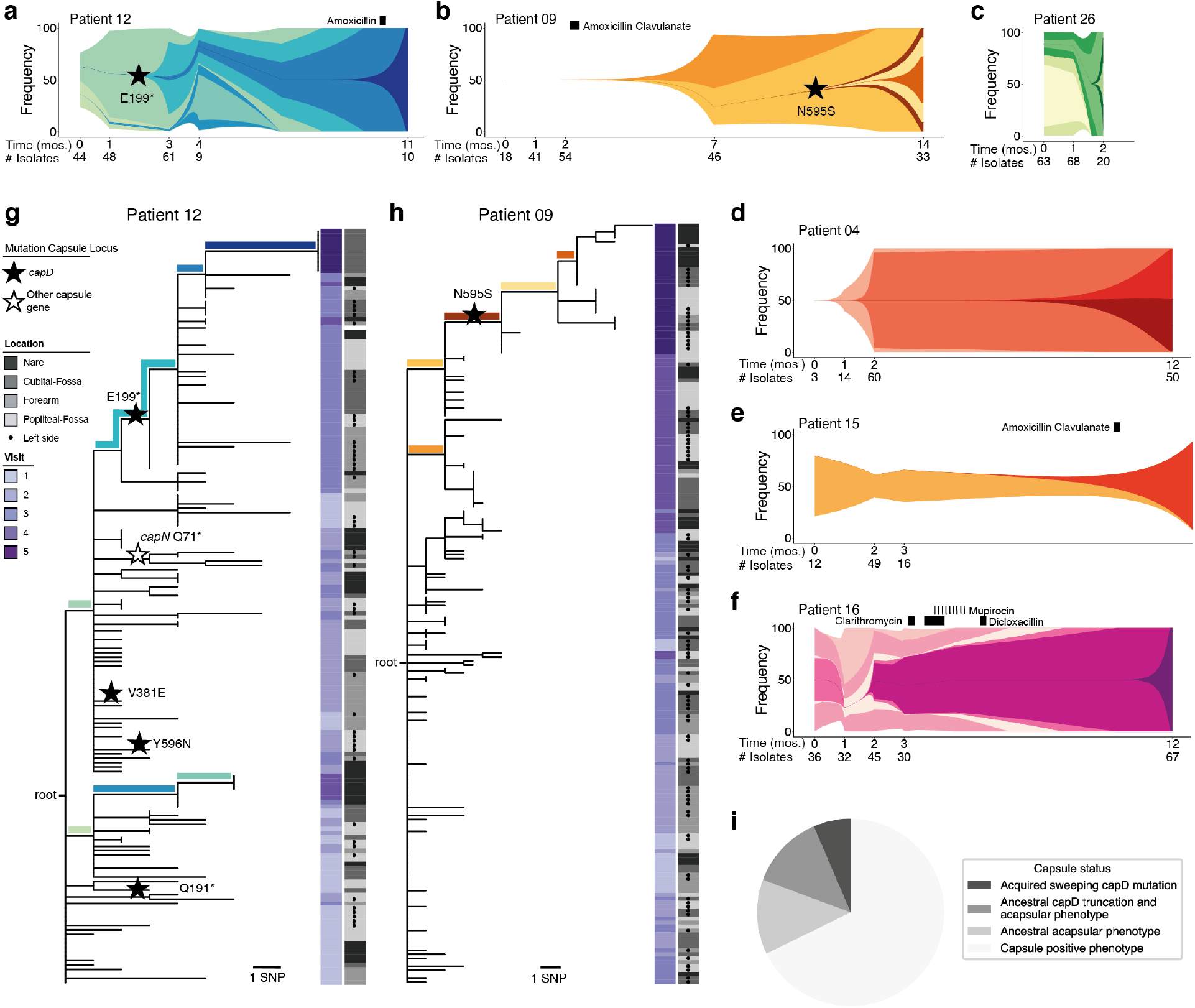
Rapid clonal spread of *de novo* mutations across the body implicates capsule loss as adaptive on treated AD skin. **a-f,** Schematics representing the on-person evolution for all patients with ≥ 99 or isolates. See **Extended Data Figures 5–6** for phylogenies of all other subjects. Each color represents a unique genotype found in at least 3 isolates and with a frequency rise of ≥ 30% for at least one visit. The shape and spacing of lines between timepoints with data is modeled based on exponential growth for maximum visibility and speculative (Methods). A star indicates a mutation in *capD*. Black rectangles indicate time and duration of antibiotic treatment. **g, h,** Maximum parsimony tree of evolution of patient 12 (g) and patient 09 (h). The location from which each isolate was taken is indicated in gray (dots indicate left side of body) and timepoint is indicated by the purple heatmap. Stars indicate a mutation within the capsule locus. Branches representing mutations defining genotypes in panels a and b are colored accordingly. **i,** Pie chart indicating the capsule status for 31 major and minor lineages recovered from all 23 patients. Sweeping capsule mutations were observed in 2 patients’ *S. aureus* populations, 4 lineages were determined genetically and by *in vitro* phenotyping to have lost the capsule prior to the start of our study, and 4 lineages were determined capsule-negative by *in vitro* phenotyping only (**Supplementary Table 3**).

The ladder-like phylogenies we observe here, showing succession of closely related genotypes, strongly support a role for on-person mutation, rather than transmission from another person or outside source (e.g. **Figure 3h**). In theory, however, these phylogenies could arise if transmission between individuals was so frequent that there was back-and-forth transmission between each pair of visits. To understand intraperson transfer, we examined the one pair of siblings in our data set. These siblings did not share mutations derived during the study period -- ruling out frequent back and forth transmission. Altogether, these analyses suggest that new *S. aureus* genotypes commonly emerge via on-person *de novo* mutation on people being treated for AD, some of which subsequently sweep the on-person population.

### Adaptive mutations alter polysaccharide capsule

The speed of genotype replacement raises the possibility that the underlying mutations provide a competitive advantage on AD skin. To test if this process was adaptive, or arose from a neutral bottleneck (e.g. growth following a reduction in cell number during treatment), we investigated mutated genes for adaptive signatures. We first searched for evidence of parallel evolution, i.e. multiple mutations in the same gene within a single patient ^37^. In patient 12, we observed four different mutations in *capD*, each in separate isolates. This mutational density is unlikely given the small number of mutations in this patient (P = 4.3 x 10^-5^, simulation), and all four mutations resulted in either amino acid replacement or a premature stop codon, further supporting an adaptive role for mutations in this gene (**Figure 3g**). The *capD* gene encodes an enzyme that performs the first step in synthesizing the capsular polysaccharide of *S. aureus*. One of the *capD* mutations in patient 12, premature stop codon E199*, is associated with a spread and replacement event in this patient (1 of 11 mutations associated with replacement in this patient). Strikingly, a spread and replacement event in patient 9 is also associated with a mutation in *capD:* a nonsynonymous N595S mutation (**Figure 3h**). While isolates containing this mutation have detectable capsule expression (Methods; **Supplementary Table 3**), this amino acid position is conserved within *S. aureus* and may be critical for native capsule structure (**Extended Data Figure 7**). These observations of parallel evolution within a patient, parallel evolution across patients, and association with replacement events suggest that alterations in *capD* provide a survival advantage for *S. aureus* on AD skin.

We did not detect comparably strong signals of adaptive *de novo* mutation across patients in *S. aureus* genes other than *capD*. Varying selective pressures across patients may have weakened adaptive signatures, and some adaptive events may have been removed by our strict filtering criteria (Putative adaptive signatures, including mutations associated with sweeps, are listed in **Supplementary Table 4**). In addition, while the presence of mobile elements varied across isolates in a lineage (**Extended Data Figure 8**; **Supplementary Table 5**), we did not find any selective sweep that was associated with the gain of a mobile element.

### Capsule loss is common in AD globally

The polysaccharide capsule of *S. aureus* has been well studied, and it is generally considered a virulence factor that shields the pathogen from phagocytosis and the innate immune system ^39–41^. However, the loss of the *S. aureus* capsule has been observed previously to emerge *in vitro* and *in vivo*^42,43^, including the epidemic USA300 clone, and it has been shown that capsule-negative *S. aureus* exhibits improved adherence to fibrinogen, platelets, and endothelial cells ^44–46^. The adaptive emergence of *capD* truncation mutations on AD skin suggests an advantage for the acapsular phenotype on AD skin.

We next sought to understand the generality of selection for *capD* alterations in our cohort. It is possible that replacement events involving *capD* emerged prior to the start of our study, given our inference of recent replacement events (**Figure 2c**). Alternatively, a patient’s *S. aureus* population may have been initially founded by a strain with a nonfunctional *capD*. We tested all patient isolates for a complete *capD* open-reading frame and observed that the major lineage of four additional patients had ancestral truncation mutations in this gene. Three of these patients share the same frameshift mutation (single base insertion at an adenine hexamere of *capD*) also found in many CC8 strains, indicating that this mutation was carried on their founding strains. The remaining lineage had a unique *capD* truncations (10 bp deletion). We also searched for capsule loss driven by non-truncation mutations or mutations in other genes by performing immunoblots, which identified two additional major lineages without detectable polysaccharide capsule *in vitro*. In total, 24% of the major lineages were acapsular at enrollment (**Figure 3i, Supplementary Table 3**).

To better understand whether loss of *capD* is generally beneficial for *S. aureus in vivo* or specifically advantageous on AD skin, we leveraged publicly available genomes of isolates from people with and without AD. We analyzed 276 *S. aureus* isolate whole-genomes from 110 AD patients, 67 healthy carriers, and 99 individuals who had other infections (blood-stream, soft tissue, bones, and joint; **Figure 4a**) ^28,47–49^. These samples were collected from individuals in Denmark, Ireland, and the UK.

**Figure 4.**
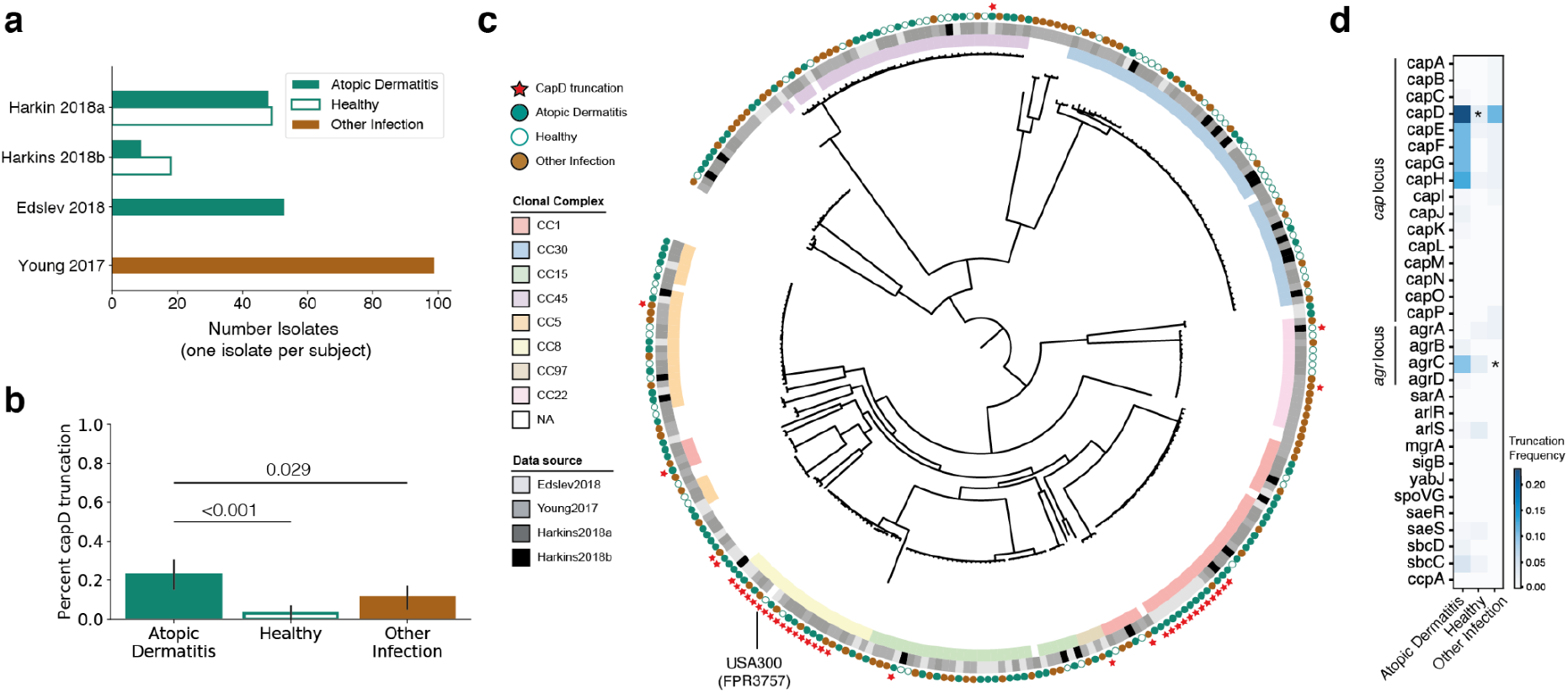
Meta-analysis of published data confirms enrichment of capsule loss on AD skin. **a,** The number of *S. aureus* genomes downloaded from each of 4 public studies are shown ^28,47–49^. Isolates were collected either from people with AD (green), healthy individuals (white), or from a variety of other diseases, including bloodstream, bone/joint and soft-tissue infections (brown). When multiple isolates were available from a subject, only the isolate with the highest number of available reads was analyzed. **b,** The fraction of isolates with a truncated or missing *capD* is plotted as a function of isolation context. The 95% binomial proportion confidence interval is shown. **c,** Phylogenetic tree showing relationship between all *S. aureus* analyzed, generated using a reference-based approach and maximum-likelihood reconstruction (Methods). Isolates are labeled with squares indicating their membership in global lineages and the study of origin, with circles indicating isolation context. Red stars indicate isolates without a full length *capD*, showing 9 different independent occurrences including two expanded clades. **d,** Analysis similar to that in (b) was performed for other genes involved in capsule regulation, and the percentage of isolates from each context lacking a full length copy of each gene is shown as a heatmap. * P < 0.001 vs. AD (one-sided Fisher’s exact test). See **Extended Data Figures 9–10** for further analysis of the capsule locus and agrC loss.

Strikingly similar to our observation that 24% of AD patients carry a strain encoding a truncated CapD, 22.5% of AD-associated isolates in the public data lack a full-length *capD* gene. This represents a significant increase relative to isolates from healthy controls (7.2%, P < 0.001; one-sided Fisher’s exact test) and from those with other types of infections (10.5%, P = 0.020) (**Figure 4b**). Phylogenetic analysis confirms several independent emergences of *capD* truncations, supporting the notion that *de novo* loss of *capD* can drive *S. aureus* adaptation (**Figure 4c**). Notably, 78% of capD-truncation isolates come from two independent, globally successful clones with recent expansions, including a CC8 clade from which USA300 is descended and a CC1 clade with a large deletion spanning *capD-capH*. Nevertheless, the observed signal remains significant when isolates from CC8 are excluded (AD vs. healthy: P < 0.001, AD vs. other infection: P = 0.016) and when the entire dataset by Edslev et al. ^48^ is removed, which included most of the CC1-*capD* negative isolates (AD vs. healthy P = 0.014). Furthermore, when considering all genes in the capsule locus, we observe an even stronger statistical enrichment of truncation mutations in isolates from AD (P < 0.001, vs. healthy; **Extended Data Figure 9c**). These genomic differences indicate that selective pressures felt by *S. aureus* on AD skin are distinct from those on healthy patients or in the context of acute infection.

To understand if capsule loss via mutations in other genes is also enriched in AD, we repeated the same analysis for genes known to be associated with capsule regulation ^50,51^. Interestingly, *agrC*, the histidine-kinase sensor in the *agr* quorum-sensing-dependent virulence regulation system, also showed a significant enrichment of independent loss-of-function mutations in AD compared to other infections (P < 0.001) but not isolates from healthy patients (**Figure 3d**; **Extended Data Figure 10**). While recent work has suggested that retention of functional *agr* is associated with the onset of AD ^27^, *agrA* loss has been previously documented on the skin of a patient with AD ^28^. Loss of the *agr* system is known to suppress capsule production, among other virulence-associated pathways ^50,52^. While the selective advantage driving frequent *agr* loss in AD could be independent of capsule, this observation is consistent with an advantage for an acapsular phenotype on AD skin.

What is the mechanistic basis of acapsular advantage on AD skin? Possible, non-mutually exclusive, selective pressures for capsule loss include: (1) immune escape from capsule-targeted antibodies or innate immune components; (2) increased adherence to AD skin via improved accessibility of surface adhesin proteins; and (3) alleviation of metabolic costs. Antibiotic resistance is unlikely to be a driver of capsule loss, as resistance-associated genomic features were generally depleted from isolates from AD context in our meta-analysis, with the exception of fusidic acid (**Extended Data Figures 10 and 11**) -- a common topical antibiotic whose efficacy we confirmed is not impacted by capsule negativity (MIC 0.05ug/mL for all strain pairs tested from Pt. 9 and Pt. 12; Methods).

Metabolic cost may explain the preponderance of mutations in *capD* -- the first gene in the biosynthetic pathway -- rather than an even distribution across all capsule-associated genes. To understand the cost of capsule production, we measured growth rates in rich media, and found that acapsular isolates grew 1.4 - 2% faster than their capsular ancestors (an advantage amplified over multiple generations; Methods; **Extended Data Figure 12**). To test if this growth advantage translated to better growth or worse disease *in vivo*, we compared the ability of these strains to cause skin injury in an mouse model that mimics an AD flare ^53,54^ and to colonize the unbroken skin of the mouse ear ^1^(**Supplementary Note**; Methods). The *in vivo* growth advantage of the acapsular strain did not translate to significantly worse disease or bacterial loads in these short-term models (< 1 week). Additionally, three AD isolates with point mutants that did not abrogate the capsule entirely did not show a growth advantage *in vitro* (**Extended Data Figure 12**). Together, these data suggest additional selective pressures for the acapsular phenotype on AD skin that were not apparent in our mouse models, possibly due to the short duration of the *in vivo* experiment. The ability of acapsular strains to evade antibodies ^55,56^ and increase adhesion ^44–46^ have been demonstrated in mouse models of other infections and may play a role during long term colonization on humans.

## Discussion

*S. aureus* is among the most successful opportunistic pathogens of humans, colonizing a third of the world’s population, responsible for numerous outbreaks in healthcare facilities, and causing a variety of acute and chronic diseases^3^. Here, we report that *S. aureus* rapidly adapts via *de novo* mutations on people being treated for AD, and that loss-of-function mutations in a *S. aureus* gene closely associated with virulence, *capD*, are frequent on AD, but not healthy, skin.

Loss of function mutations in *capD* are present in global lineages, yet new bacterial genotypes with mutations in this gene also arise on individual patients with AD, spread across a patient’s skin, and replace their ancestors within months. Our finding that loss of function in the capsule pathway is associated with AD is surprising in the lack of a consistent signal in AD-association at the level of sequence types, clonal complexes, or other phylogenetic groupings. The discrepancy across scales of genomic units raises the possibility that ecological dynamics (e.g. priority effects^57^) or selective tradeoffs limit the specialization ability of *S. aureus* to AD skin ^58^.

At a global scale, it is possible that the advantage of being acapsular on AD skin may translate to all skin and may in part explain the success of the acapsular *S. aureus* USA300 lineage in causing skin infections and spreading to epidemic levels outside the hospital^42^; future studies are needed to test whether acapsular mutants have an increased capacity for survival on unaffected spread and inter-person spread.

We cautiously propose that small molecules or other therapies that specifically target *S. aureus* strains without a functional capsule locus might be combined with selective forces on the host to lower the burden of *S. aureus* colonization in AD. Future work further characterizing the mechanistic basis behind the CapD-negative advantage on AD skin will be critical to the design of such therapies, as unexpected consequences may emerge from selection for the capsular phenotype, which is more virulent in other disease contexts^39,59^.

This study focused on *S. aureus* genotypes that could be easily recovered from swabs at 7 body sites. We did not seek to characterize community members at low abundance, and it is possible that additional minority lineages and genotypes were present at undetected levels or at additional locations on the body (**Extended Data Figure 13**). Similar limitations exist for culture-independent approaches, as it is difficult to distinguish between sample cross-contamination and low abundance colonization^60^. Our data indeed support a model in which lineages can persistently colonize despite lack of detection; we recover the exact same lineages from four patients after visits at which no *S. aureus* was recovered (**Figure 1b**). Notwithstanding, we find no cases of reemergence of replaced genotypes, supporting the durability of observed intralineage sweeps on the skin of people treated with AD. As all patients were treated for AD, additional work is needed to test if *S. aureus* has similar dynamics in untreated AD and to understand the specific contribution of treatment, and its resultant decrease in population size ^61^, on *S. aureus* on-person evolutionary dynamics.

Together, our results highlight the potential of *de novo* mutations for altering bacterial competitiveness in microbiomes, highlight the power of mutation tracking for identifying new potential therapeutic directions, and suggest that whole-genome resolution may be required for predicting the impact of microbial strains on complex diseases.

## Methods

### Study cohort and sample collection

Patients were recruited from the Dermatology Clinic at the National Institute for Pediatrics in Mexico City under a protocol approved by the Institutional Review Boards of the NIP (042/2016) and Massachusetts Institute of Technology. Inclusion criteria for enrollment were: ages 5 to 18 years, diagnosis of AD according to modified Hanifin and Rajka criteria ^62^ and SCORAD ≥ 25 at first visit. We preferentially enrolled patients that had not taken topical or systemic antibiotics for the past month, though one patient (P15) did take antibiotics immediately prior to enrollment. Patients were sampled at up to five timepoints (Figure 1A). Some deviation from this schedule occurred due to patients’ schedules, as well as the disruption caused by the earthquake in September 2017. Patients were counseled avoidance of irritants and adequate use of fragrance-free moisturizers and/or emollients 3 times daily. Treatment with anti-inflammatory medication (either class II-VII topical corticosteroids or the calcineurin inhibitor tacrolimus 0.1%) was prescribed considering multiple factors including skin sites affected, severity of AD, and age of the patient. Patients were instructed to apply the topical corticosteroids 2x or 3x daily on affected sites until improvement was achieved. Once improved, patients were instructed to continue using the topical medication twice weekly on previously affected areas for two more weeks. At each clinical visit patients were re-evaluated and either instructed to stop the topical medication, change the type/class of medication, or to continue using the same medication twice daily or twice weekly based on the assessment. Some patients were additionally instructed to use dilute bleach baths (0.005%) twice weekly. Detailed information about patient care was collected at each of the first 3 follow-up visits, but not at the 6 month follow up; patients continued treatment at their own discretion between the 4^th^ and 5^th^ visits. Overview of patient data, including treatment, is in **Supplementary Table 1**.

Skin swabs were collected from seven different locations at each visit (**Figure 1A**) in Liquid Amies transport media. Swabs were directly inoculated on mannitol salt and blood agar plates and cultured for 24h at 37°C. For each culture plate, up to 10 colonies suspected to be *S. aureus* by colony morphology were selected. These colonies were restreaked on ¼ of a blood agar plate and cultured for 24h at 37°C to obtain sufficient material for DNA extraction. DNA was extracted using the Wizard^®^ Genomic DNA Purification Kit (Promega Corporation) for Gram-positive bacteria.

Dual-barcoded DNA libraries were constructed using the plexWell library prep system (SeqWell) for most samples and a modified version of the Nextera protocol for a small subset^63^. Libraries were sequenced on the Illumina NextSeq 500 using paired end 75bp reads to average of 1.3M read pairs per isolate. Demultiplexed reads were trimmed and filtered using cutadapt v1.18 ^64^ and sickle-trim v1.13 ^65^ (pe -q 20 -l 50).

We note that sequenced *S. aureus* genomes from each patient is an imperfect measure of *S. aureus* absolute abundance. In some cases, with heavy growth, it was difficult to isolate 10 colonies. In addition, some sequenced colonies were determined not to be *S. aureus* from genomic data and excluded from the analysis and some cultured isolates did not yield sequenceable libraries due to a failure in DNA extraction or library prep -- and thus could not be confirmed to be *S. aureus*. Regardless, we find good correlation between final colony count data and relative abundance inferred from 16S rRNA sequencing (**Extended Data Figure 2**).

### Assignment of isolates to lineages

Reads were aligned using bowtie2 v2.2.6 (-X 2000 --no-mixed --dovetail) against methicillin-resistant *Staphylococcus aureus* USA300-FPR3757 (RefSeq NC_007793). Candidate single nucleotide variants were called using samtools (v1.5) mpileup (-q30 -x -s -O -d3000), bcftools call (-c), and bcftools view (-v snps -q .75) ^66,67^. For each candidate variant, information for all reads aligning to that position (e.g. base call, quality, coverage), across all samples, were aggregated into a data structure for local filtering and analysis. Isolates were removed from analysis if they had a mean coverage 7 or below across variant positions (208 isolates removed of an initial 1,735). To identify isolates with contamination that would inhibit reliable read calling, the frequency of the second highest allele at each position, in each sample, was calculated (minor allele frequency) and those isolates with a mean minor allele frequency above 0.03 across candidate sites were removed (21 isolates). We filtered candidate SNVs using a publicly available protocol (see Data Availability) similar to that previously published^37^, with the following filters. Basecalls were marked as ambiguous if the FQ score produced by samtools was above −30, the coverage per strand was below 2, or the major allele frequency was below 0.85. Variant positions were filtered if 5% or more of all isolates were called as ambiguous (this removes most of the ‘accessory’ genome) or if the median coverage across strains was below 3, or if no unmasked polymorphisms remained. These filters retained 60,973 SNVs across 1,506 isolates. A maximum-likelihood tree was built with RAxML v8.2.12 ^68^ using the GTRCAT model, with rate heterogeneity disabled (-V). Sporadic isolates that were phylogenetically confined within the diversity of another patient were likely mislabeled and removed (7 isolates removed, leaving a final total of 1,499 isolates). The phylogenetic tree was visualized with iTol ^69^. For each isolate the sequence type and clonal complex was inferred using SRST2 ^70^ and the *S. aureus* MLST/CC database was obtained March 2020 (with 5,964 sequence types)^71^. All 22 isolates from patient 17 were manually assigned to CC30 after failing SRST2 clonal complex assignment, despite matched allele types for 6 out of 7 alleles that define CC30; the CC30 assignment is also supported by the phylogeny (Figure 1c). Isolates that failed sequence typing were ignored for CC-dependent analysis and are not shown in Figure 1b (70 out of 1,499). Lineages are defined as phylogenetically closely related *S. aureus* from the same patient which differ by less than 100 SNVs on the core genome **(Extended Data Figure 1)**. Analysis of minor lineage detection dependency on the number of isolates sampled at a given visit suggests that deeper sampling would have been unlikely to detect additional lineages from most patients (**Extended Data Figure 2c**). A second phylogeny performed using the methicillin-sensitive *S. aureus* FDA209P (NCTC 7447, RefSeq NZ_AP014942, NZ_AP014943) genome as a reference for alignment ^72^ resulted in the same lineage groupings (data not shown).

### Within-patient phylogenetic reconstruction using *de novo* assemblies

We constructed patient-specific pan-isolate assemblies in order to capture true genome-wide diversity; this approach can capture the accessory genome only found in a single isolate ^14^. For patients colonized by multiple lineages, we only analyzed isolates from the major lineage. Only isolates with ≥ 80% reads assigned to *S. aureus* on the species level (assigned using kraken2, default parameters, and standard database build Sep 24 2018) ^73^ were included for *de-novo* assembly, leaving 1,215 isolates. For each lineage, we concatenated up to 250,000 reads from each member isolate and assembled a reference genome with SPAdes ^74^ (v3.13, careful mode). All scaffolds of size 500b or larger were annotated using prokka ^75^ (v1.14.6), which was supplied with a list of proteins (--*proteins*) from nine publicly available *S. aureus* genomes and six accompanying plasmids (NCBI accession: NC_002745, NC_003140, NC_002758, NC_002774, NC_009641, NC_003923, NC_002952, NC_002953, NC_005951, NC_002951, NC_006629, NC_007795, NC_007793, NC_007790, NC_007791, NC_007792). The assemblies for each patient are summarized in **Supplementary Table 6**.

We aligned the data of all initial 1,704 isolates from major lineages to their respective patient-specific assemblies using bowtie2 ^76^ using the same parameters as described for *assignment of isolates to lineages* analysis. Isolates and candidate single nucleotide variants were processed with identical methods as listed in *assignment of isolates to lineages*, but with the following modifications: (1) To remove isolates with an inflated mean coverage across the assembled genomes due to high-copy number plasmids we required that at least 10% of the genome covered at 8X; (2) To detect variations within the accessory genome, the allowed maximum fraction of ambiguous isolates per site was increased from 5% to 25%; (3) To remove variants that emerged from recombination or other complex events, we identified SNVs that were less then 500b apart and covaried highly across isolates within a patient (a SNV was considered to highly covary with another SNV if its covariance was in the 90% percentile across covariances calculated with the focal SNV); these positions were removed from downstream analysis. (4) To remove cross-contaminated samples, isolates were omitted if they had minor mean allele-frequency of 9% across verified SNVs (removing 6 isolates of patient 19). Overall, these filters removed 284 isolates, leaving 1,422 isolates and 915 *de novo* on-person SNVs across 23 subjects (**Supplementary Table 7**). Phylogenetic reconstruction was done using *dnapars* from PHYLIP v3.69 ^77^. Each lineage’s tree was rooted using the isolate with the highest coverage from the most closely related lineage as an outgroup (based on *assignment of isolates to lineages* analysis).

### Molecular clock and TMRCA

We estimated the number of *de novo* mutations per isolate using all positions variable within a patient lineage. The ancestral state of each variant is defined by the allele called in the lineage-specific outgroup (the isolate with the highest coverage from the most closely related lineage; see above). If this was not available, the major allele across the set of isolates used as an outgroup for any lineage was used. For Patient 26, we checked the predicted ancestral genotype using *treetime* ^78^ after noticing a significantly elevated molecular clock (using procedures below); we changed the ancestral allele of the outgroup at four basal mutation positions to match the best molecular clock fit predicted by *treetime* ^78^. We normalized the number of mutations per isolate to the number of positions on the patient-specific assembled reference with a depth of at least 8X. We inferred the molecular rate using linear regression implemented in scipy v.1.3.1 ^79^ and the 95% confidence interval using the two-sided inverse Students t-distribution. Time to the most recent common ancestor (tMRCA) was calculated for each patient data from each patient’s earliest visit with at least 10 isolates. For each of the six highly colonized patients the tMRCA was calculated using their inferred mutation rate, while for all other patients the median molecular rate of the six highly colonized individuals was used (2.41 × 10^-6^ substitutions/site/year).

### Signatures of within-person adaptive evolution

Each within-patient dataset was searched for genes with either of two signatures of adaptive evolution: parallel evolution at the gene level ^11^ and hard selective sweeps ^80–82^. First, we identified cases when two or more mutations arose in a single gene within a patient, with a minimum mutation density of 1 mutation per 1000 bp. We calculated a p-value for enrichment of mutations in each gene using a Poisson distribution^83^. Only the *capD* gene in patient 12 had a significant p-value after Bonferroni correction for the number of genes on the genome (**Supplementary Table 4**). Second, we searched for genomic positions at which the mutant allele frequency rose by at least 30% between visits. For each SNV, we assigned the ancestral allele as the allele found in the patient-specific outgroup, or, if that was not available, we used the allele present in the patient-specific assembly. We compared lists of genes with candidate adaptive signatures across patients using CD-HIT (v4.7, 95% identity)^84^. All candidate signatures of adaptive evolution are reported in **Supplementary Table 4**.

### Creation of Muller plots

We visualized the change in frequency of mutations observed on each patient with ≥99 isolates using Mueller plots. Patient 04 and patient 15 yielded only two isolates at visit 4 and visit 2, respectively; these timepoints were omitted from visualization. We only drew mutations with an observed variation in frequency of ≥0.3 between timepoints. We converted mutation frequencies into genotype trajectories using *lolipop* v0.8.0 (https://github.com/cdeitrick/Lolipop), SNVs were grouped into a single genotype (color) if derived allele frequency difference was less than 8%. To make successive sweeps more visible, we used a custom-made python function to generate intermediate genotype frequencies between sampling timepoints. In brief, we created 100 time units per month, assumed mutations swept sequentially, and applied an exponential growth or decline following the function of frequency x = ε^((ln(f_1_ - ln(f_0_)) / g), with f_0_ is the frequency at preceding visit or 0, f_1_ is the frequency at the next visit and g is the number of time units (see code availability). The extended table was used again as the input for *lolipop*, which provided the input tables for plotting using the R package *ggmueller* v0.5.5 ^85^.

### Mobile genetic element analysis

We identified genetic gain and/or loss events based on the depth of coverage of each isolate aligned to its patient-specific assemblies along the six individuals with at least 99 isolates ^14^. To avoid spurious results due to uneven coverage across samples or genomic regions, we calculated a z-score for each position by normalizing each isolate’s depth of coverage at each position by sample and by position. We identified genomic regions greater than or equal to 5,000 bp long, for which each position’s z-score was below a threshold of −0.5 with a median threshold below −1.0. Identified candidate regions were further filtered to have at least one isolate with a median coverage of 0 and at least one with an average depth of at least 10X. These cutoffs were determined using an interactive python module for visualization of coverage across the entire candidate region across all samples (**Supplementary Table 5**).

### Capsule typing

*S. aureus* capsule serotyping was performed as described previously ^86^. Briefly, duplicate tryptic soy agar plates were spot inoculated in a grid pattern with *S. aureus* isolates and incubated overnight at 37°C. The colonies were blotted onto nitrocellulose filter membranes for 5 min at ambient temperature. Adherent colonies were fixed to the membranes by heating them at 60°C for 15 min. After two washes with 10 mM sodium phosphate buffer-0.85% sodium chloride (phosphate-buffered saline; PBS) to remove excess cells, the filters were immersed in a solution of trypsin (1 mg/ml; Sigma) for 60 min at 37°C to remove protein A from the bacterial cells. After two washes in PBS, the filters were blocked with 0.05% skim milk for 1 h and washed in PBS containing 0.05% Tween 20 (PBST). Each filter was incubated with capsule type-specific rabbit polyclonal antisera (diluted 1:4,000 in PBST) at 37°C for 1 h. After washing in PBST, horseradish peroxidase-conjugated donkey anti-rabbit immunoglobulin (diluted 1:5000) was incubated with each filter for 1 h at 37°C. After three washes, KPL 1-Component TMB Membrane Peroxidase Substrate was added to the filters. A purple color developed within 5 min and was scored visually from 0 to 4+. The reactivities of the clinical isolates were evaluated by comparison with those of control *S. aureus* strains (serotypes 5, 8, and nontypeable isolates) included on each filter. Results are presented in **Supplementary Table 3**.

### Growth rate

Overnight cultures were inoculated into fresh TSB at a 1:100 dilution. Cultures were grown at 37°C with shaking, for 24h in a Tecan plate reader (Infinite F Nano+, Tecan Trading AG, Switzerland). Absorbance at 595 nm was measured every 5 minutes. The plate containing growing cultures was sealed using its own lid. The growth rate was calculated using a regression on log-normalized data and selected for a single time interval across all strains where the r^2^ of the estimated growth rate was > 0.999. Code and raw data available at: github.com/vedomics/growthrateR.

All experiments were performed with representative test isolates carrying *de novo* mutations in the *capD* gene from patients enrolled in this study. For each test isolate, a phylogenetically closely related and basal control isolate was chosen. Multiple test isolates were used in cases where all available isolates contained 1-2 additional mutations compared to the most closely related basal isolate observed (see **Supplementary Table 7)**.

### Fusidic Acid Susceptibility

Overnight cultures of patient isolates of *S. aureus* were diluted and plated on varying concentrations of fusidic acid (Sigma-Aldrich), ranging from 0.02ug/Ll to 2.5ug/mL. Both capsule positive and capsule negative strains were found to be susceptible at 0.05ug/mL, although the emergence of resistant mutants was common on both backgrounds.

### Public *S. aureus* isolate sequencing data analysis

We investigated publicly available data to verify if loss of CapD is associated with *S. aureus* colonizing AD skin. We obtained whole-genome sequence data from 4 different publications analyzing *S. aureus* in healthy individuals, AD patients, or individuals with other *S. aureus* infections ^28,47–49^. When data from multiple isolates per patient was available, the isolate from the site of infection (if available) with the highest coverage was used. To examine gene content, we performed *de novo* assembly for each isolate using SPAdes (v3.13, --careful)^74^ and annotated the assembly using Prokka (v1.14.6)^75^ as described above. Isolates with assembly lengths of < 2.6M nucleotides were not considered and removed from consideration (removing 6 isolates). After this removal, we analyzed 276 isolates from 276 patients. To understand the relationship between isolates, we performed an alignment-based phylogenetic reconstruction using *S.aureus* COL (sequence version number: NC_002951.2, NC_006629.2) and the same filters as above. In addition, we included USA300-FPR3757 (RefSeq NC_007793) as a sample, by simulating raw reads for input to our cross-lineage phylogenetic analysis (cutting its genome in segments of size 150 b in steps of 1 b to simulate reads). The maximum likelihood phylogeny was built using RAxML v8.2.12 (-m GTRCAT)^68^ and visualized with iTol ^69^.

We inferred the ORF status for all capsule and capsule-associated genes using the annotated assemblies and BLAST+ v2.7.1 ^87^. Details about each query gene are available in **Supplementary Table 8**. We compared the best BLAST match to each query gene for overlap with an annotated reading frame. We accepted an ORF as complete if the start and end of the best BLAST hit were each within 100 bp of a gene start or end of a gene in the annotated assembly. *S. aureus* genomes colonizing humans are known to carry 1 of 2 predominant *cap* loci (type 5 or 8)^39^ and 1 of 4 *agr* types ^88^ containing homologous versions of the same genes; for these cassettes we performed analysis only for the respective loci with the best BLAST match to a given isolate. Results are reported in **Figure 4**, **Extended Data Figures 9–10** and summarized in **Supplementary Table 9**. When only data from publications reporting both AD and controls are included, the enrichment of *capD* premature stop codons in AD vs controls remains significant (P=0.026).

### Collectors Curve Analysis

For each patient’s sampling point (visit) with at least 10 isolates a collector curve was created for dMRCA or identified SNVs (**Extended Data Figure 13**). For each patient-visit dataset we re-sampled with replacement 0 < x < n isolates each x sampled 100 times, calculated the within-sample diversity, and plotted the mean and standard deviation.

## Author contributions

M.T.G-R. and T.D.L designed the clinical cohort. M.T.G-R. enrolled patients and collected clinical samples. C.R-G. cultured bacteria from clinical samples and extracted DNA. T.C.L. and T.D.L. prepared genomic libraries. F.M.K. and T.D.L performed genomic analysis and interpretation. J.C.L. performed capsule assays and provided *S. aureus* strains. V.K. and L.D. performed *in vitro* experiments. V.K., K.J.B, L.D., and I.M.C. performed mouse experiments. F.M.K. and T.D.L wrote the manuscript with feedback from all authors.

## Acknowledgements

We thank Mariana Matus and Eric Alm for assistance in the design of the clinical cohort, the BioMicroCenter at MIT for performing genomic sequencing, Samantha Choi for technical support on animal experiments, Paul F. Koffi for performing *in vitro* capsule immunoblots, Chris Manusco and Calen Mendall for experimental assistance and members of the Lieberman lab for valuable advice and feedback on the manuscript. We acknowledge support from MISTI Global Seed Funds (to T.D.L. and M.T.G-R.), the National Institutes of Health (DP2-GM140922 to T.D.L., R01AI30019 to I.M.C.), Burroughs Wellcome Fund (to I.M.C.), the Mexican Government Ministry of Taxes Program E022 for Health Research and Technological Development 2018 (to M.T.G-R.), and DFG research fellowship (KE 2408/1-1 to F.M.K.).

## Data availability

All raw data is available at SRA under BioProject identifier: PRJNA715375, PRJNA715649, and PRJNA816913. Metadata of each isolate including SRR identifier is available in **Supplementary Table 10**.

## Code availability

All code needed to reproduce the results of this study, including snakemake pipelines, are available here: https://github.com/keyfm/aureus_ad.

## Extended Data Figures

**Extended Data Figure 1.**
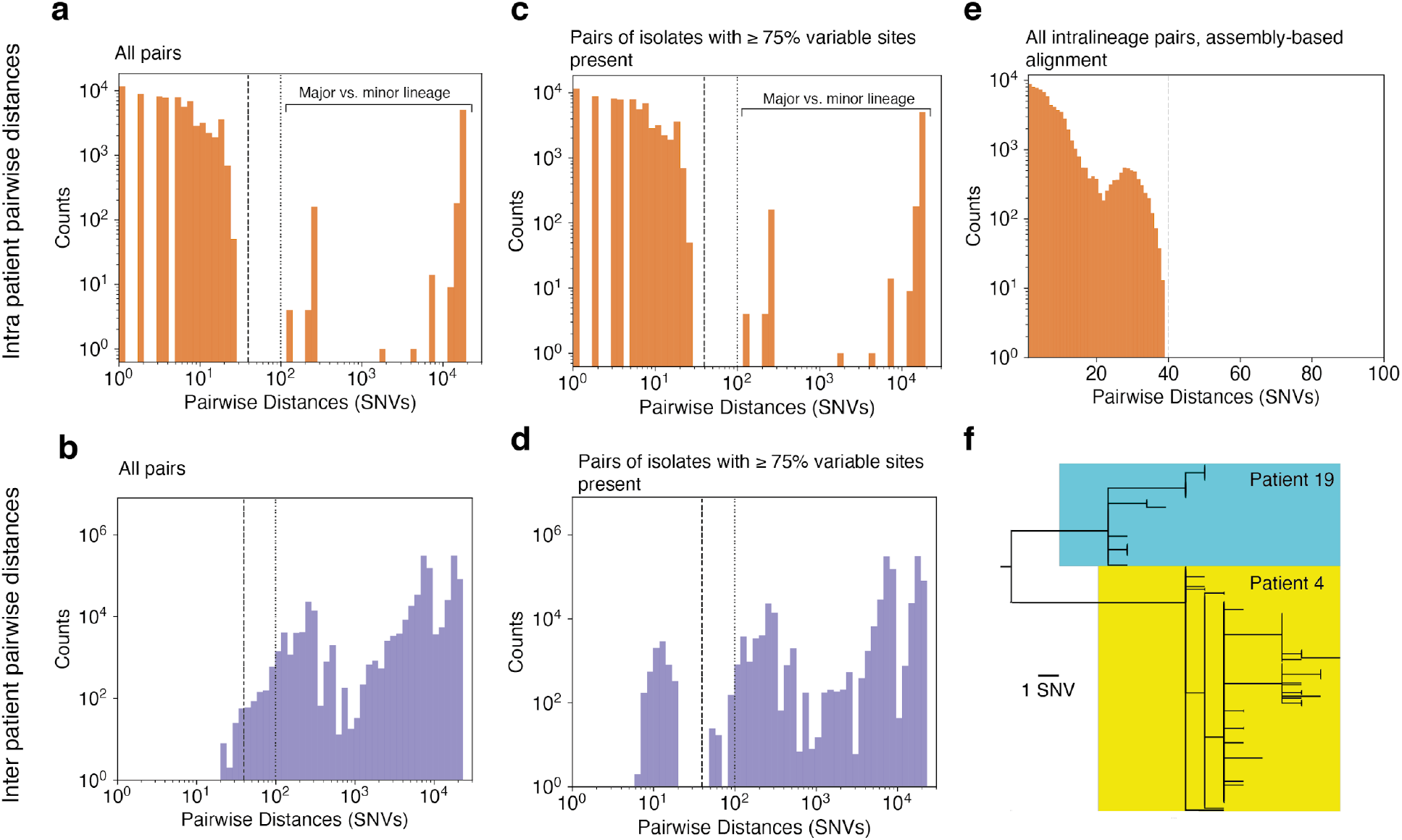
Whole-genome pairwise distances between isolates of the same patient, or between patients, enables differentiation of lineages. **a.** Histogram of pairwise distances between isolates within a patient. Isolates from the same lineage have fewer than 30 SNV differences across the core genome using an alignment-based approach (reference genome: USA300-FPR3757). Both axes are on log scale to make inter-lineage pairs more visible. **b.** Histogram of pairwise distances between isolates of different patients. We note that some pairs of isolates from different people were also separated by less < 100 SNVs (vertical dotted bar) across their core genomes using an alignment-based approach (reference genome: USA300-FPR3757); however these pairs included isolates with relatively low coverage. **c,d.** When isolates are only included in this analysis if they have at least 75% (45,742 SNVs) of all identified variable sites with coverage of 8X using an alignment-based approach (reference genome: USA300-FPR3757), a clearer separation between intra-patient and inter-patient diversity is seen at 40 SNVs (vertical dashed bar). Both axes are on a log scale to enhance visibility. **e.** Histogram of pairwise differences observed among isolates of major lineages aligned to their respective assembly-based reference genome. The Y axis is on a log scale to enhance visibility. **f.** All intra-subject pairs of isolates with a pairwise distance of < 40 SNVs in **d** are from the one sibling pair in our data set. A joint phylogeny of these subjects suggests recent seeding of their populations from a common source, or inter-patient transmission. See **Extended Data Figure 5a** and **Extended Data Figure 6** for more details.

**Extended Data Figure 2.**
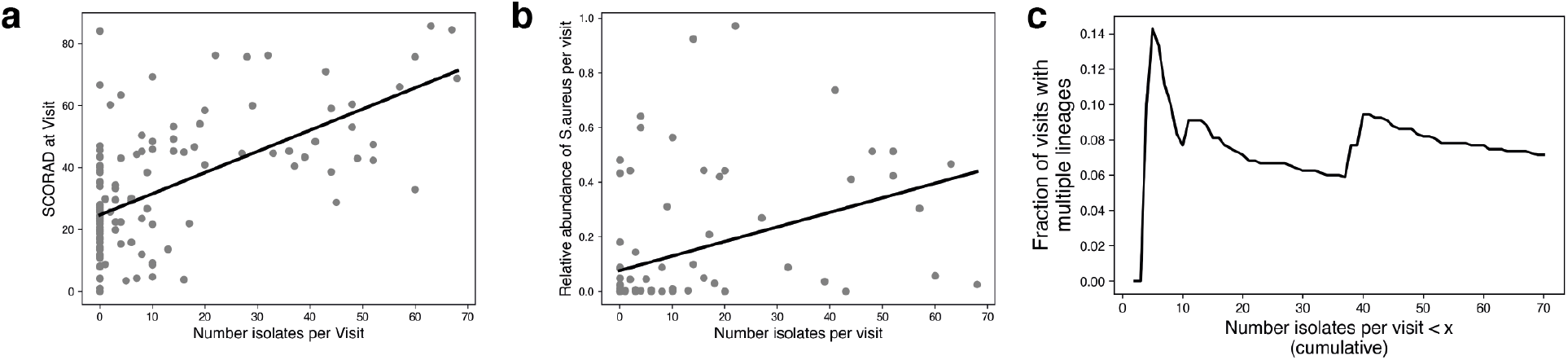
Number of recovered isolates correlates with disease severity and relative *S. aureus* abundance per visit but not with the detection of multiple lineages. **a.** Linear least-squares regression for the number of recovered isolates and disease severity (SCORAD) per visit (r^2^=0.36) demonstrates a relationship between disease severity and the number of isolates recovered. **b.** Linear least-square regression for the relative *S. aureus* frequency (16s rRNA; averaged across sites) and number of recovered *S. aureus* isolates per visit demonstrates a relationship between the relative abundance of *S. aureus* and the number of isolates recovered (r^2^=0.16). The 16S rRNA data was originally described in Khadka et al.^23^. **c.** Fraction of visits where more than 1 lineage was detected on a patient demonstrates a lack of relationship between detection of minor lineage and number of isolates cultured. Also see **Extended Data Figure 12**.

**Extended Data Figure 3.**
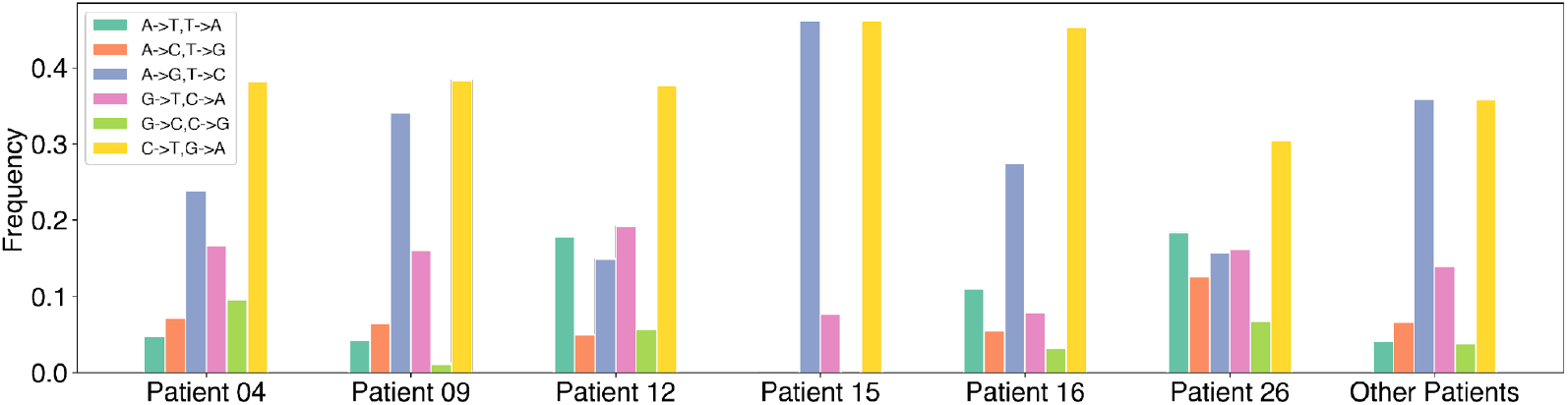
Similar mutational spectra are observed across all densely colonized patients. Mutational direction (ancestral to derived nucleotide) is based on the respective outgroup assigned to that patient (see Methods). Notably, Patient 26 does not show a mutational spectrum that deviates significantly from that of other patients.

**Extended Data Figure 4.**
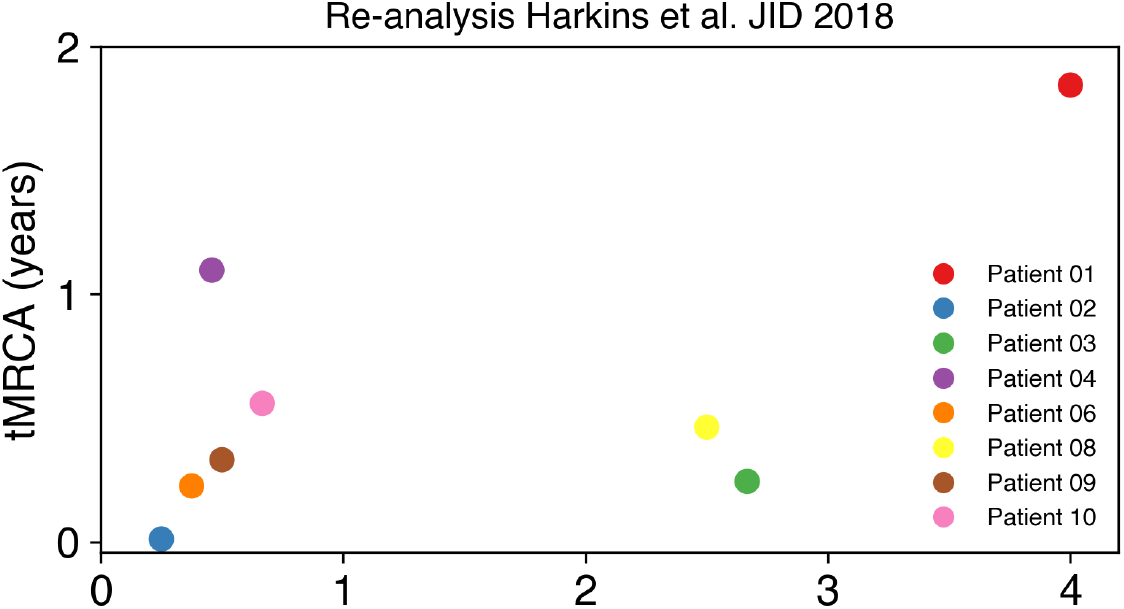
Inferred time to the most recent common ancestors across *S. aureus* isolates from AD patients presented in an independent study. Harkins et al.^28^ reported non-longitudinal *S. aureus* isolates sampled on the skin and nares from young children (0-4 years old) in the UK. We recalculated tMRCA for each individual with at least 10 isolates reported, using the same methods described for within-subject phylogenetic inference in this study--including alignment to a patient-specific assemblies and removal of putative recombination events. For each patient, we used the same outgroup as described in Harkins et al ^28^ Table S5. For positions where the outgroup remained undefined, the most frequent allele across all outgroups was used. For the molecular clock we used the median clock rate observed among the 6 densely and longitudinally colonized patients presented in this study (2.36 × 10^-6^ substitutions/site/year).

**Extended Data Figure 5.**
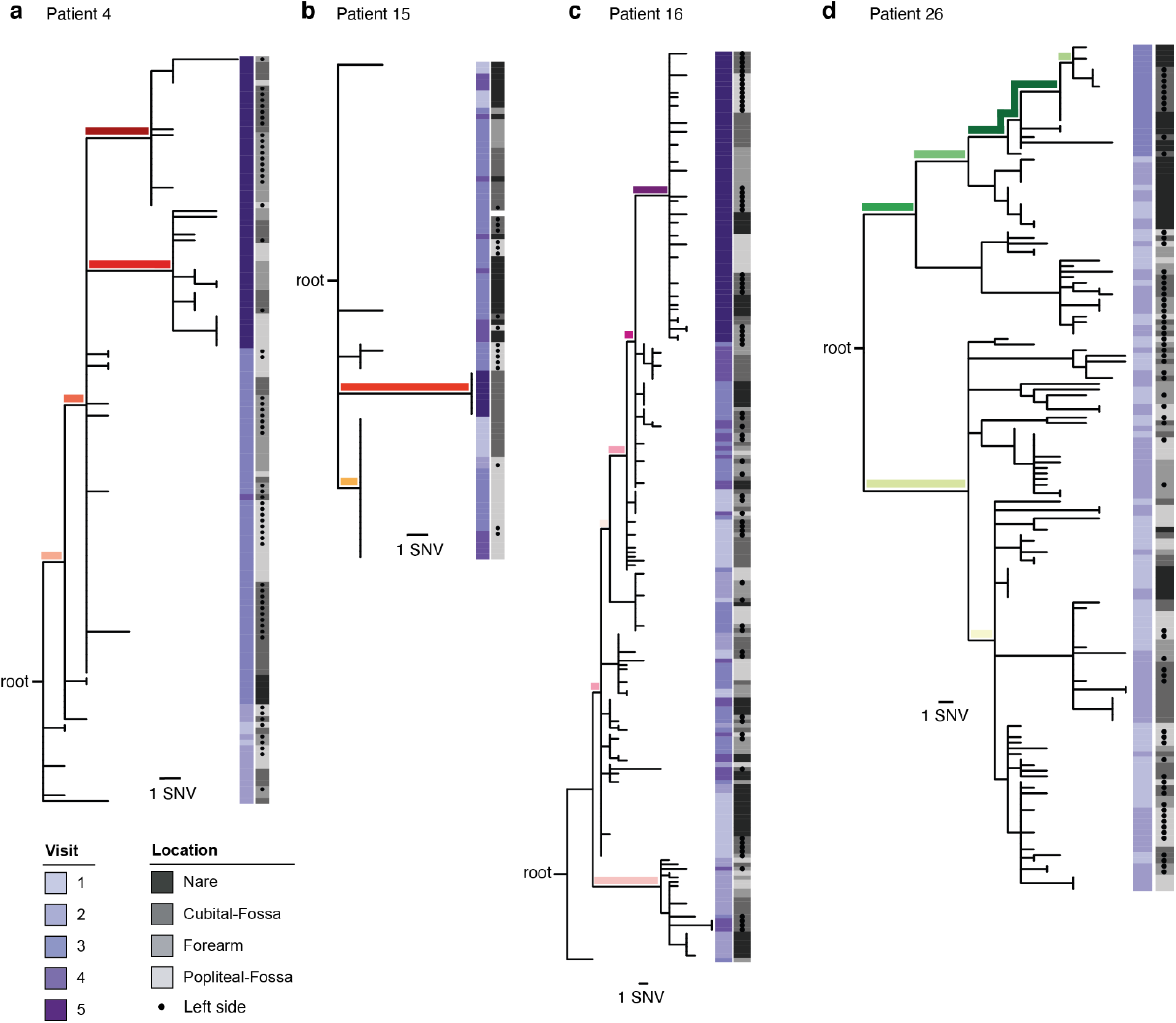
Within-patient evolutionary reconstruction of *S. aureus* for other highly colonized patients. Within-person evolutionary trees for Patients 04 (**a**), 15 (**b**), 16 (**c**) and 26 (**d**), constructed as in **Figure 3c-f.** The location from which each isolate was taken is indicated in gray (dots indicate left side of body) and timepoint is indicated with the purple heatmaps. Branches with >30% change in frequency between timepoints are colored in accordance with **Figure 3c-f.**

**Extended Data Figure 6.**
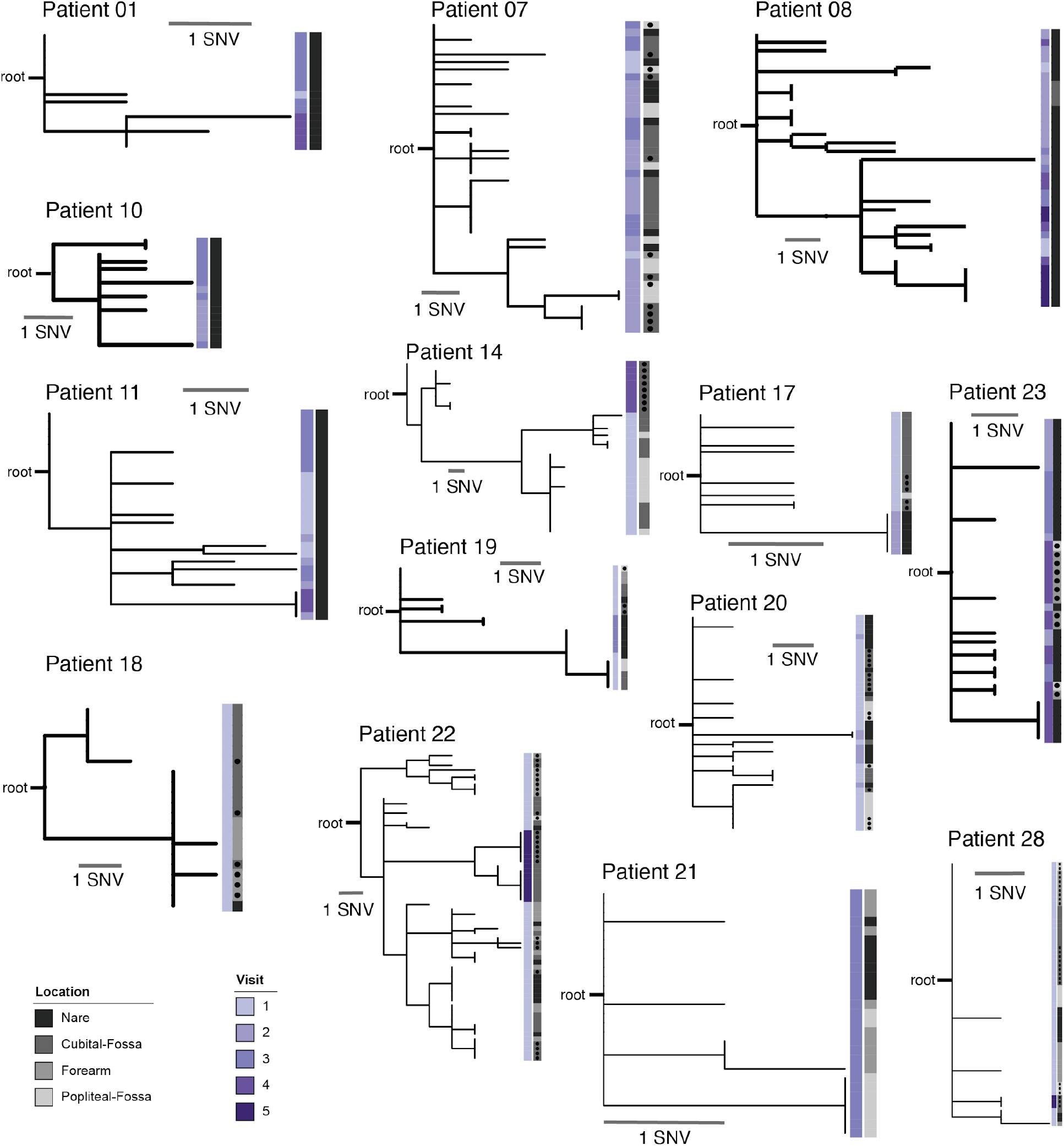
Within-patient evolutionary reconstruction of *S. aureus* genomes recovered from patients with less than 99 isolates. The patient-ID, root, and branch length is indicated for each phylogeny. Sampling visit is shown in shades of purple and sampling location of where an isolate was collected is shown in shades of gray (dot indicates left side of body).

**Extended Data Figure 7.**
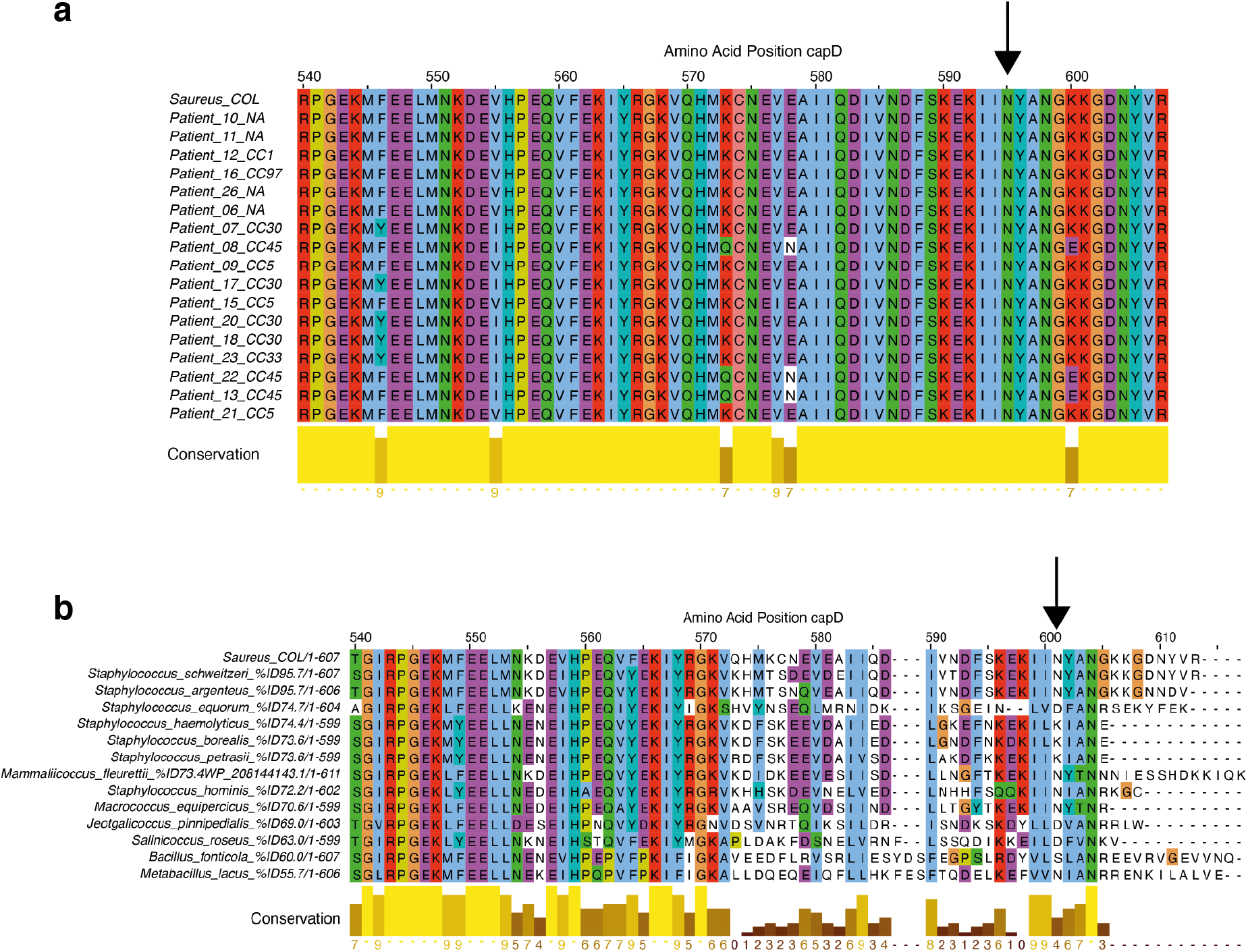
Multiple sequence alignment of the terminal *capD* amino acid sequence demonstrates conservation of amino acid position 595 (arrow) within *S. aureus* but not between species. **a.** Multiple sequence alignment of the 607 amino acid long *capD* sequence within *Staphylococcus aureus* including the COL type strain and all *capD* sequences annotated by prokka ^75^ within the patient-specific pan-genome assemblies. Assigned clonal complex shown in row label (if none ‘NA’). **b.** Multiple sequence alignment of *capD* across different species identified using BLAST ^87^. Percent sequence identity (%ID) and aligned amino acid position is shown in each row label. Amino-acid position 595 (mutated in Pt. 09 N>S) indicated by an arrow. The multiple sequence alignments have been generated using Muscle ^89^ and jalview ^90^.

**Extended Data Figure 8.**
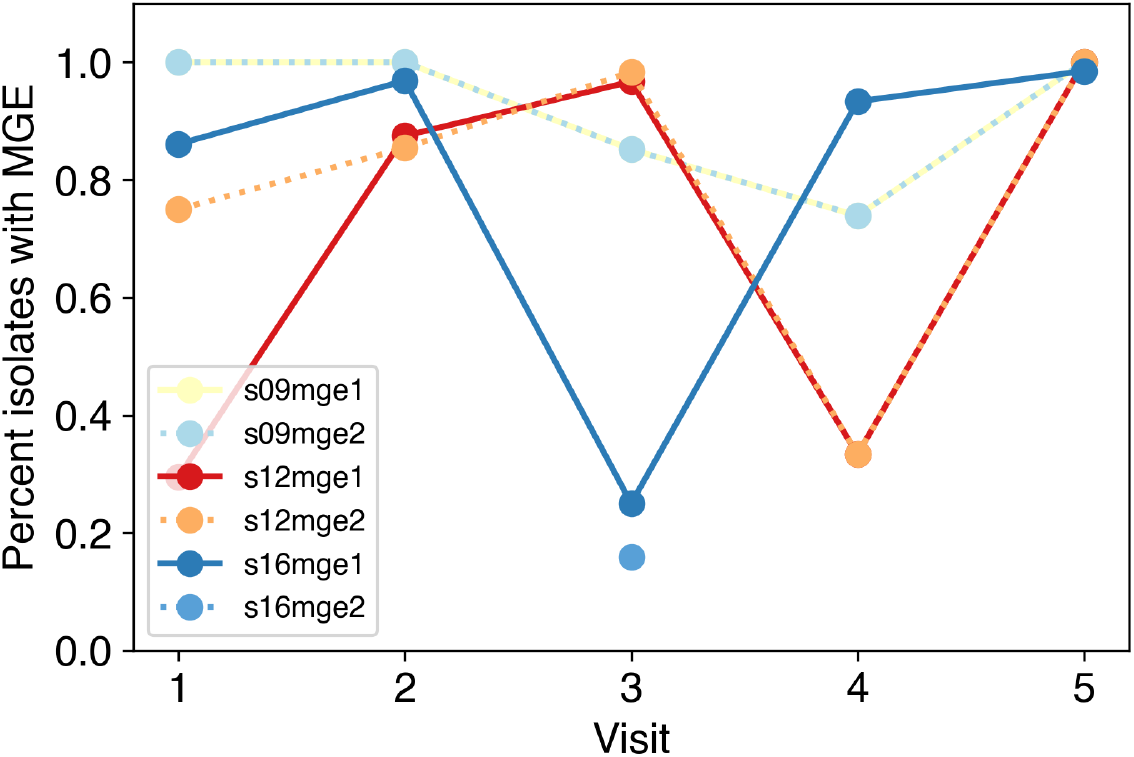
Frequency of identified mobile genetic element (MGE) among isolates per visit. Isolates were considered to harbor the MGE if the mean depth across the MGE was at least half of the mean genome-wide depth. To avoid artifacts, MGEs are reported only if at least two isolates had evidence of coverage and at least two isolates had evidence of absence. Except for an MGE detected only in Pt-16 at Visit 3, all other MGEs were present at the first sampling visit and are present in all isolates at the last visit in the same patient. Each MGE is named based on its patient of origin (e.g. S09mge1 is the first MGE from Pt-9). Gene annotations in each MGE are reported in **Supplementary Table 5**.

**Extended Data Figure 9.**
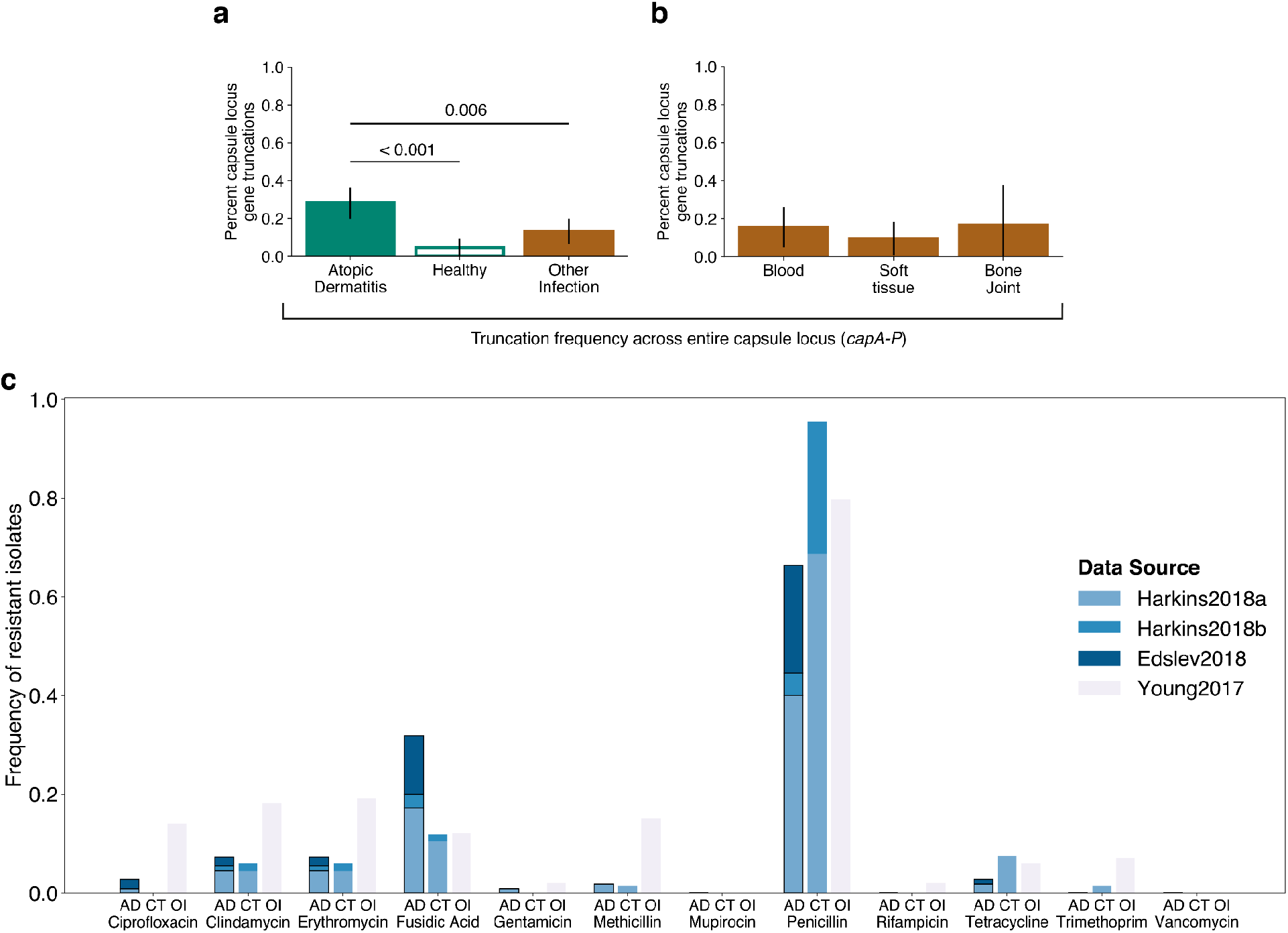
Enrichment of truncations extend across the entire capsule locus in AD patients and antimicrobial resistance profile. **a.** Frequency of truncations in genes part of the entire capsule locus (*capA-P*) for isolates derived from individuals with atopic dermatitis, healthy, and other infections. **b.** Same as (a) but only individuals with other infections are shown and differentiated by type of infection. No statistically significant difference is observed. **c.** Frequency of antimicrobial resistance inferred in the meta-analysis of the 276 isolates from 276 individuals. Mykrobe ^91^ was used to predict antibiotic resistance profiles from the publicly available genomes analyzed in **Figure 4**. The percent of isolates predicted to be resistant to the displayed antibiotics are shown for isolates derived from individuals with atopic dermatitis (AD), healthy (CT), and other infections (OI).

**Extended Data Figure 10.**
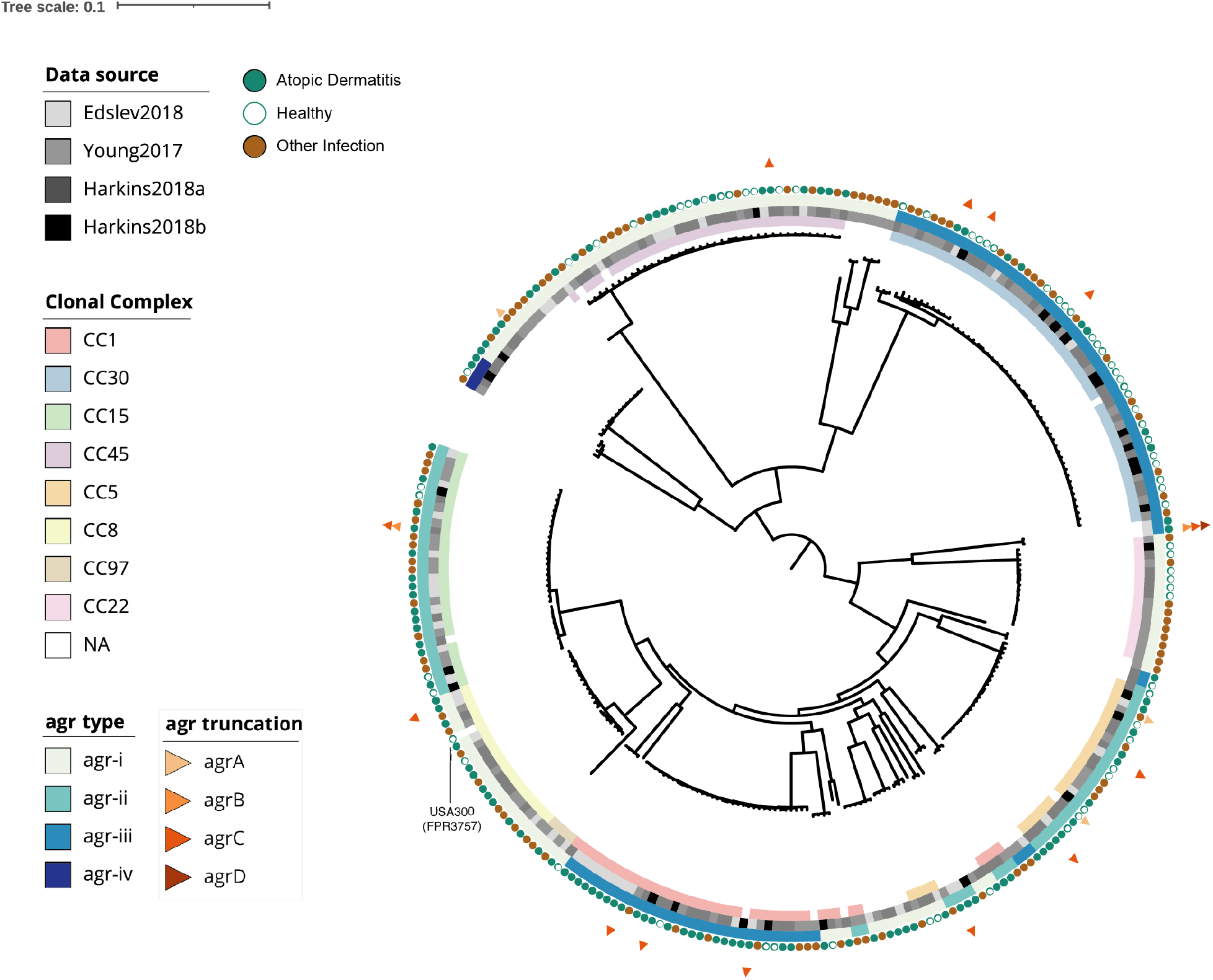
Maximum-likelihood phylogeny showing relationship for all 276 *S. aureus* isolates and strain FPR3757, using a reference-based approach (similar to **Figure 3c**). Isolates are labeled with squares indicating their membership in global lineages, the study of origin, and *agr* type, with circles indicating isolation context. Triangles indicate isolates with truncated *agr A*, *B*, *C*, or/and *D*, showing 16 different independent events.

**Extended Data Figure 11.**
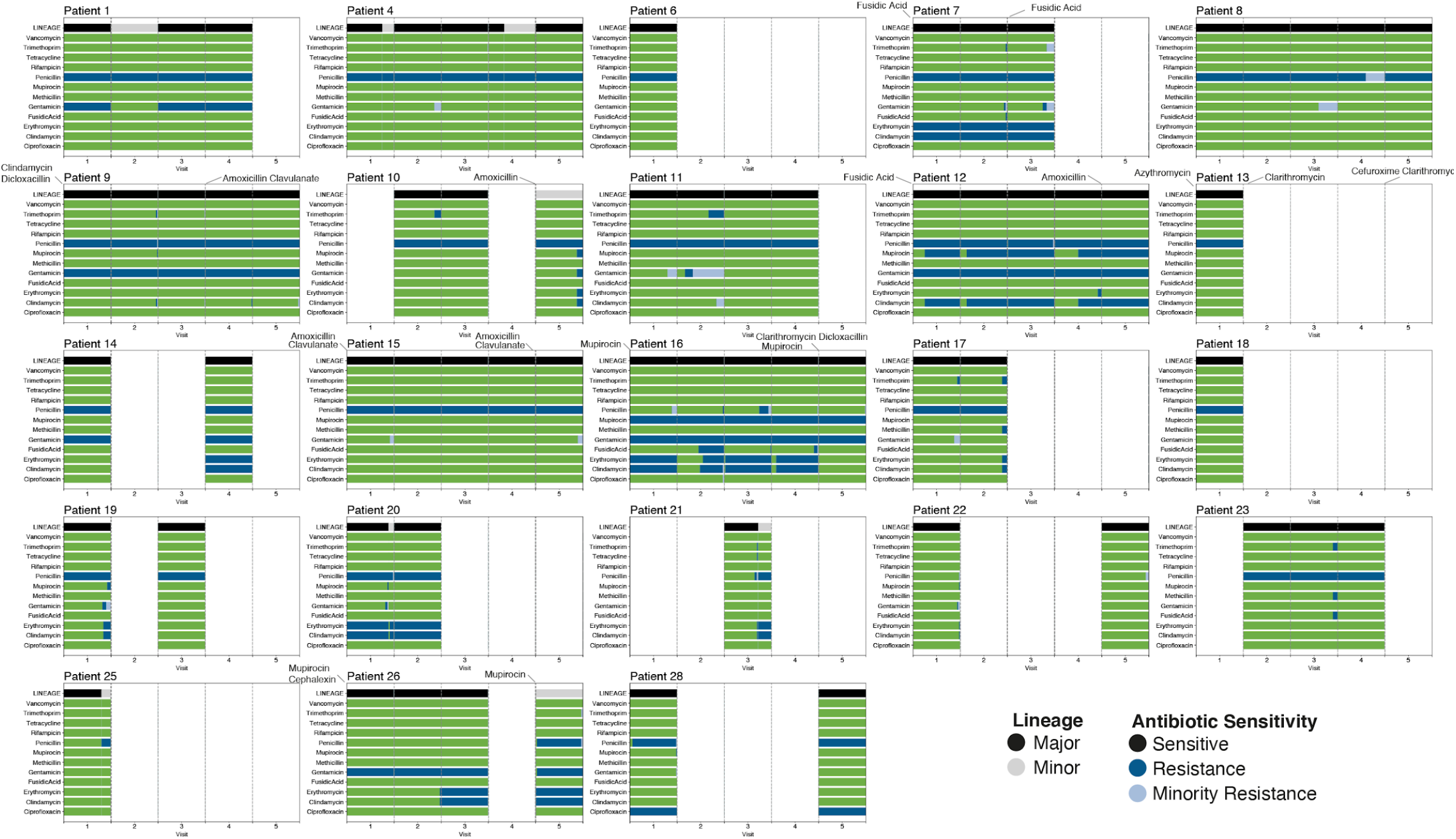
Antimicrobial sensitivity predictions for all isolates of all patients. Mykrobe ^91^ was used to predict antibiotic resistance profiles from all 1,499 isolates analyzed here. For each patient and visit, the percent of isolates from the major on-person lineage (black) and minor on-person lineage (gray) is shown. The percent of isolates predicted to be antibiotic resistant is shown in blue; location along the x-axis is arranged such that any vertical line within a patient reflects the same lineage isolate (i.e. resistance profiles of co-colonizing major and minor lineage can be compared). Administration of antibiotics is indicated on top of plot (see also **Supplementary Table 1**).

**Extended Data Figure 12.**
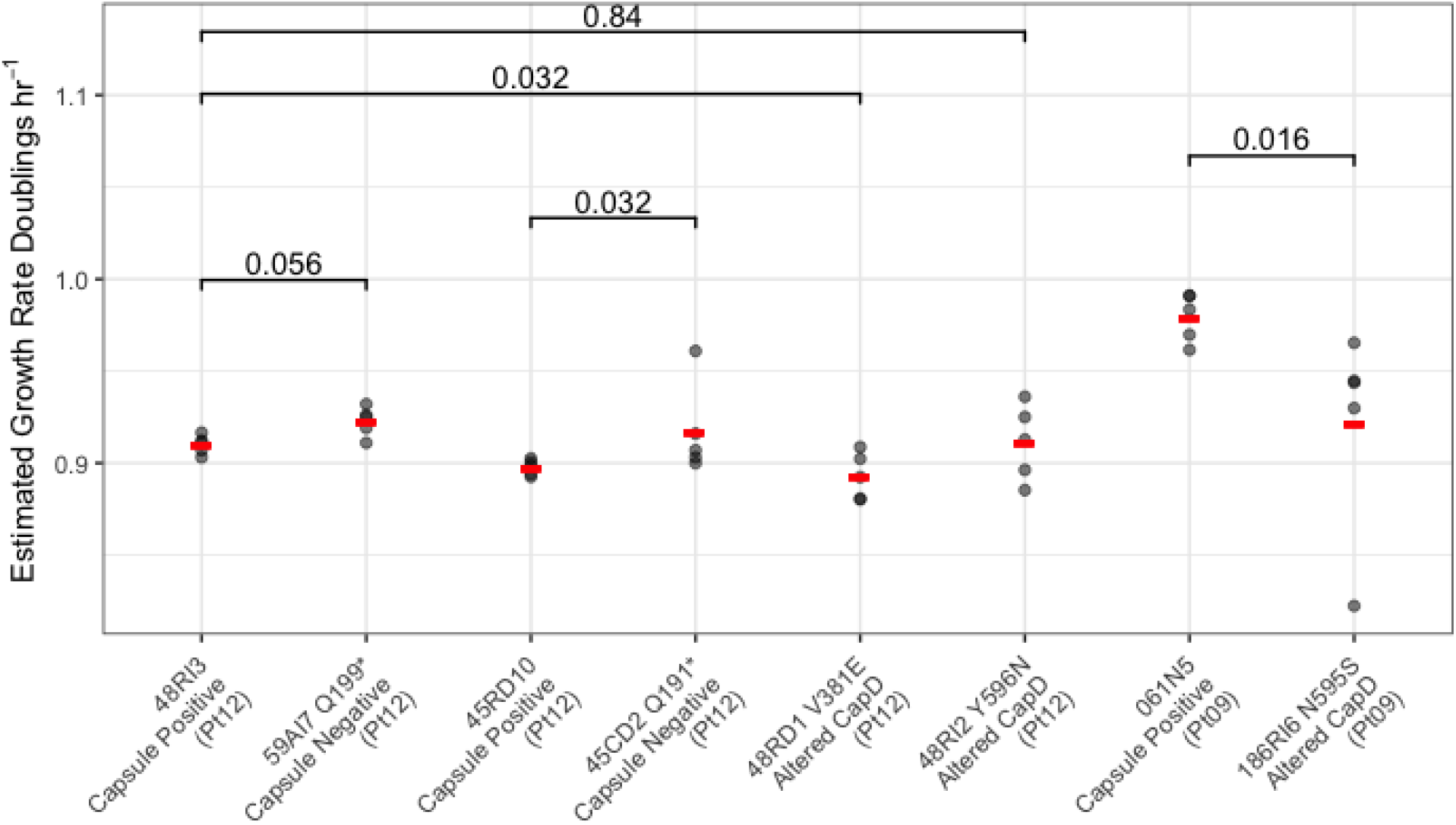
Growth advantage of acapsular versus capsular isolates. Growth rates in TSB of isolates carrying *de novo* mutations in the *capD* gene as well as closely related phylogenetically basal strains (Methods), showing a growth advantage for the acapsular isolates. P-values of Wilcoxon rank sum tests are shown. Strain 59-AI7 shows a 1.4% increase in growth rate over 48-RI3, and strain 45-CD2 shows a 2% growth rate increase over 45-RD10. Each capsule mutant was compared to the most closely related isolate in our collection, as indicated by the lines and p-values at top. Some pairs had 1-2 additional mutations outside the capsule locus, see **Supplementary Table 7.** Red lines indicate means.

**Extended Data Figure 13.**
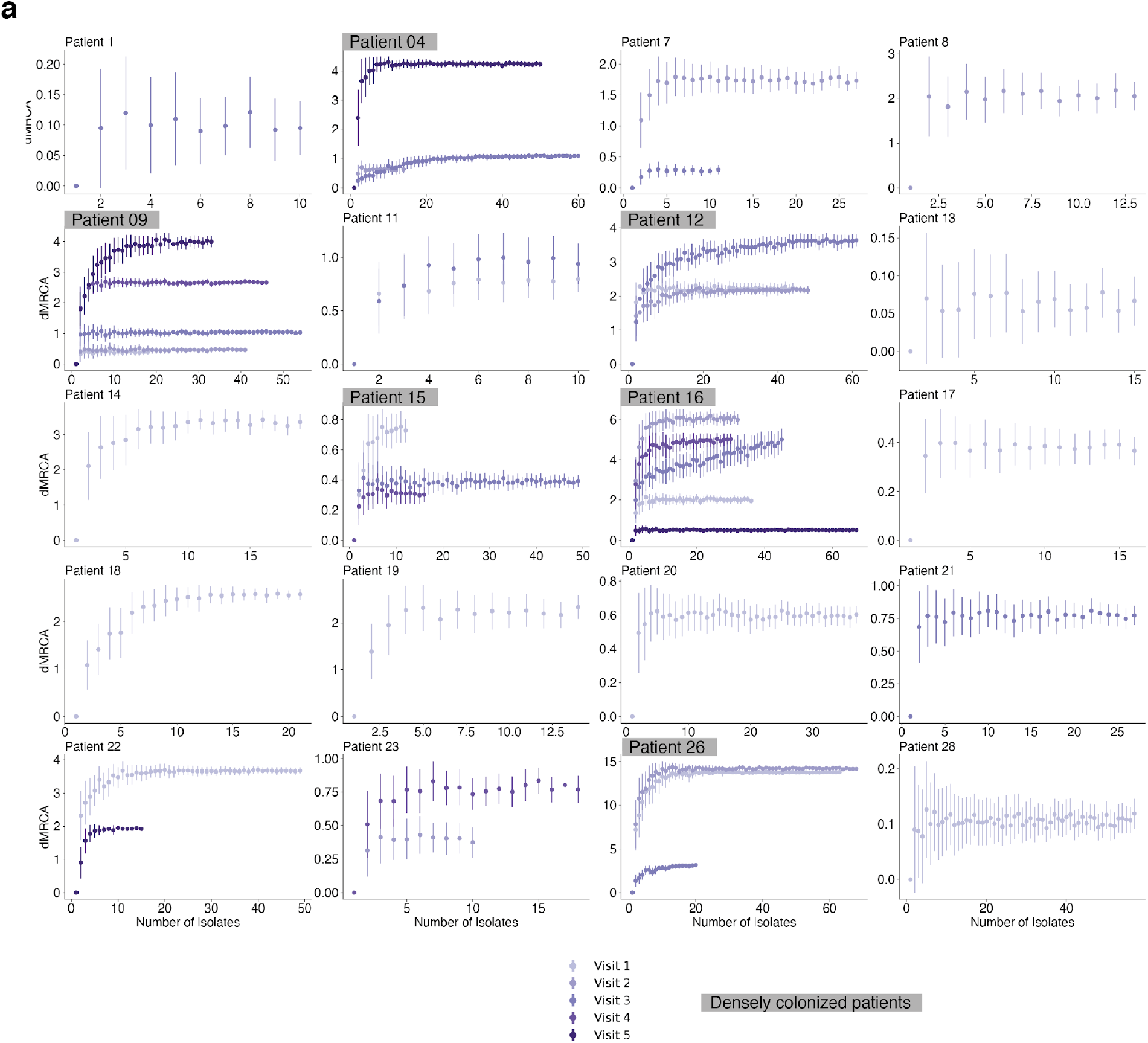
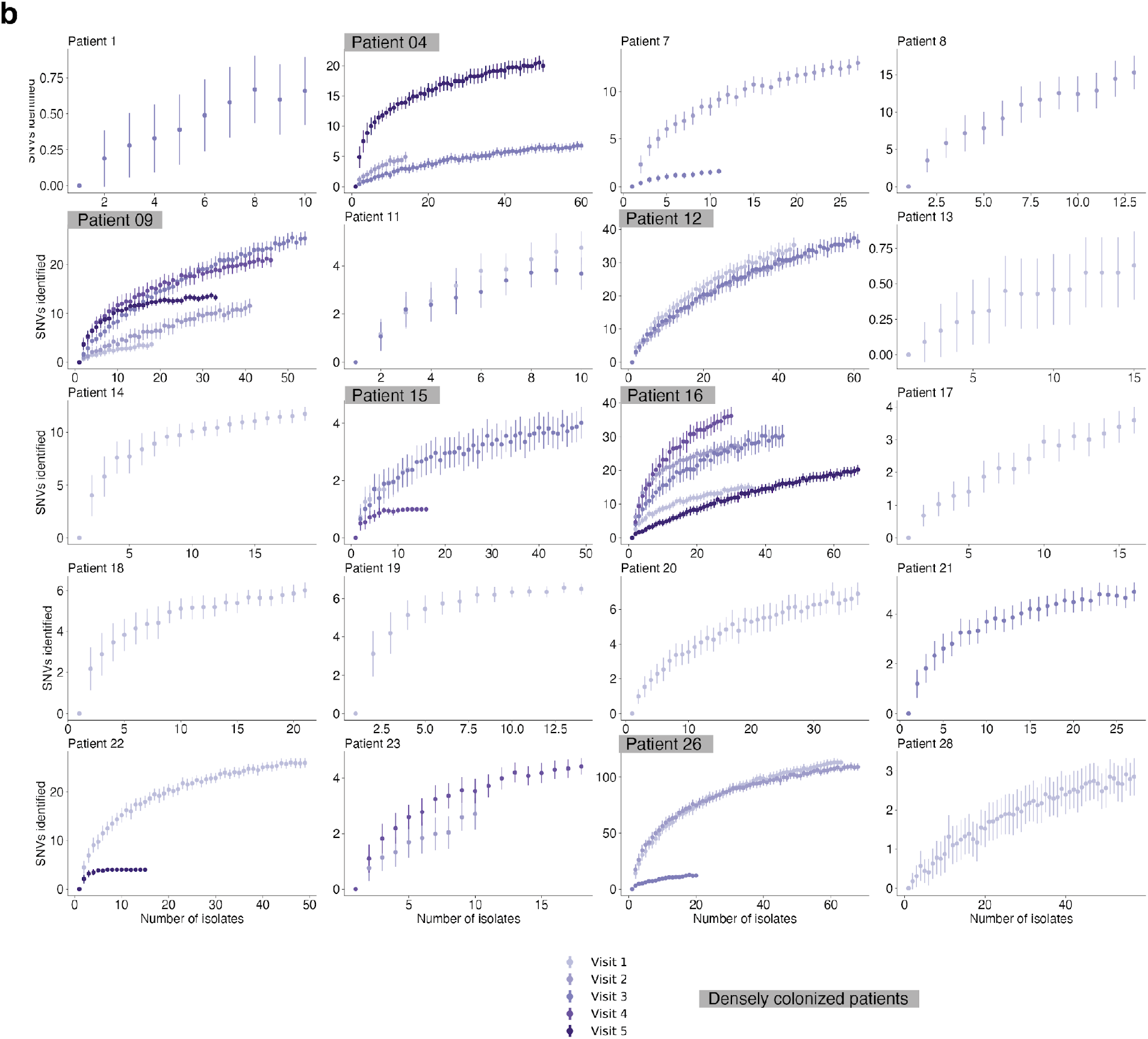
Collector curves demonstrate sufficient sampling of patients intra-lineage diversity but not rare newly emerged SNVs. For each sampling point (visit) with at least 10 isolates a collector curve for (**a**) dMRCA or (**b**) identified SNVs was created, combined on a single plot per patient. For each patient-visit dataset we re-sampled with replacement 0 < x < n isolates each x sampled 100 times, calculated the within-sample diversity, and plotted the mean and standard deviation. The dMRCA results suggest sufficient sampling for most patients, whereas the SNV results suggest that rare variation remains, likely reflecting newly emerged SNVs.

## Supplementary Note

### Previous *in vivo* experiments with laboratory strains

In a previous version of this manuscript available on bioRxiv ^92^, we reported a difference between Newman WT and a Newman Cap5D deletion mutant in an epicutaneous infection model in mice. However, the two strains were found to have additional point mutations between them, including a nonsynonymous mutation in the sensor kinase *saeS* that confounded the reported impact of cap5D deletion on disease severity in this model. Here, we report only on experiments performed on strains isolated from patients, which do not demonstrate strong effect of capsule loss in mouse and tissue culture models (**Supplementary Figure 1** and **2**). Future studies are needed to show if the advantage of the acapsular phenotype *in vivo* is potentially triggered by the adaptive immune system or something on skin from people with atopic dermatitis not captured in our models.

### Methods for Mouse Experiments

#### Bacterial Strains and Cultures for *in vivo* experiments

All experiments were performed with representative bacterial isolates carrying *de novo* mutations in the *capD* gene from patients enrolled in this study. For each test isolate, a phylogenetically closely related and basal control isolate was chosen. Sometimes the test isolate contains additional mutations compared to the most closely related basal isolate observed.

*Patient 12:*

Control: 048-RI3
Test: 059-AI7 (Q199* in *capD;* synonymous mutation in a hypothetical protein)

*Patient 9:*

Control: 061-N5
Test: 186-RI6 (N595S in *capD*; G278D ComE operon protein 3 *comEC*; intergenic mutation between two hypothetical proteins)

Bacteria were grown to stationary phase overnight at 37°C in Tryptic Soy Broth (TSB) at 250 r.p.m. Stationary phase cultures were diluted 1:100 in fresh TSB and grown for 3.5 hours to approximate mid-log phase. Cultures were then centrifuged at 800xg for 15 minutes, and the resulting pellet washed twice in phosphate buffered saline (PBS). Cells were resuspended to a concentration of ~10^10^ CFU/mL. Serial dilutions were plated on Tryptic Soy Agar (TSA) plates to confirm inoculum cell densities.

#### Animals

Eight-week old C57BL/6 female mice from Jackson Laboratories (Bar Harbor, ME, USA) were housed in specific pathogen free animal barrier facilities at MIT in individually ventilated isolator cages under a 12 h light/dark cycle with ad libitum food and water access. Euthanasia was performed by CO_2_ inhalation.

All animal experiments were approved by the Institutional Animal Care and Use Committee (IACUC) at MIT.

#### Epicutaneous Skin Infection and Bacterial Load Measurements

Two days prior to epicutaneous infection, mice were shaved and the remaining hair removed using depilatory cream (Nair) along the length of their back/flank. A sterile 1 cm^2^ square of gauze was soaked in 100 uL of prepared *S. aureus* inoculum (~10^9^ CFU/mL for each strain, see crosshatches in figures) and applied to the flank skin. The gauze was secured using bio-occlusive film (Tegaderm, 3M) and the tape checked and repaired daily to maintain its integrity. Gauze soaked in sterile PBS was used as a control. Seven days post-inoculation, mice were euthanized by CO_2_ and the dressing removed. Skin under the gauze was immediately scored for disease severity according to the following criteria: oedema (0-3), erythema (0-3), skin scale (0-3), and skin thickness (0-3) and totaled for a max skin score of “12.”. A higher score indicated more severe disease/inflammation. For bacterial load, the flank skin immediately under the gauze (~1 cm^2^) was excised and resuspended in 1 mL cold PBS contained in a 2 mL microcentrifuge tube. The tissue was cut into smaller pieces using sterilized scissors, two metal ball bearings (4.5 mm) were added to each tube, and the tissue was homogenized using a TissueLyser II (Qiagen, Germany) at 25 s^-1^ for 5 min. Homogenates were briefly spun down, serially diluted in PBS, and then plated for CFU counts. Bacterial identity was confirmed by plating on ChromAgar Colorex *S. aureus* (ChromAgar, France) plates to differentiate any native microbiota that may have been present. A total of two replicates were performed. The analysis of all animal experiments was blinded.

For colonization experiments, bacteria were prepared as above and ~10^7^ CFU were applied to each ear in a 50 uL volume. Mice were euthanized 24 h later and tissue was harvested, homogenized, and serially diluted in PBS. Bacterial identity was confirmed by plating on ChromAgar Colorex *S. aureus* plates.

#### Tissue culture and intra-epithelial invasion assay

The CCD 1106 KERTr cell line (ATCC CRL-2309), comprising human epithelial keratinocytes, was seeded into flat-bottom 24-well plates in Keratinocyte Serum Free Media (Gibco) with added Keratinocytes Supplements including bovine pituitary extract and human recombinant epidermal growth factor (Gibco) and further supplemented with additional 35 ng/mL human recombinant epidermal growth factor (BD). Cells were grown at 37 C in 5% C02 until confluent. Confluent cells were washed and infected with *S. aureus* isolate strains 48-RI3, 59-AI7, 61-N5, 186-RI6. After two hours of incubation, cells were washed and cell media containing 40 ug/mL Lysostaphin (Fisher Scientific) was added and the mixture incubated for an additional hour. All strains tested were equally susceptible to killing by lysostaphin. The epithelial cells were then washed with PBS and detached by incubation with 100 uL of 0.25% Trypsin (Sigma) for 5 minutes at 37 C with 5% C02 and lysed with 400 uL of 0.1% Triton-X. Cell lysates were diluted and plated on TSA to enumerate the number of intracellular *Staphylococci*.

## Supplementary Figures

**Supplementary Figure 1.**
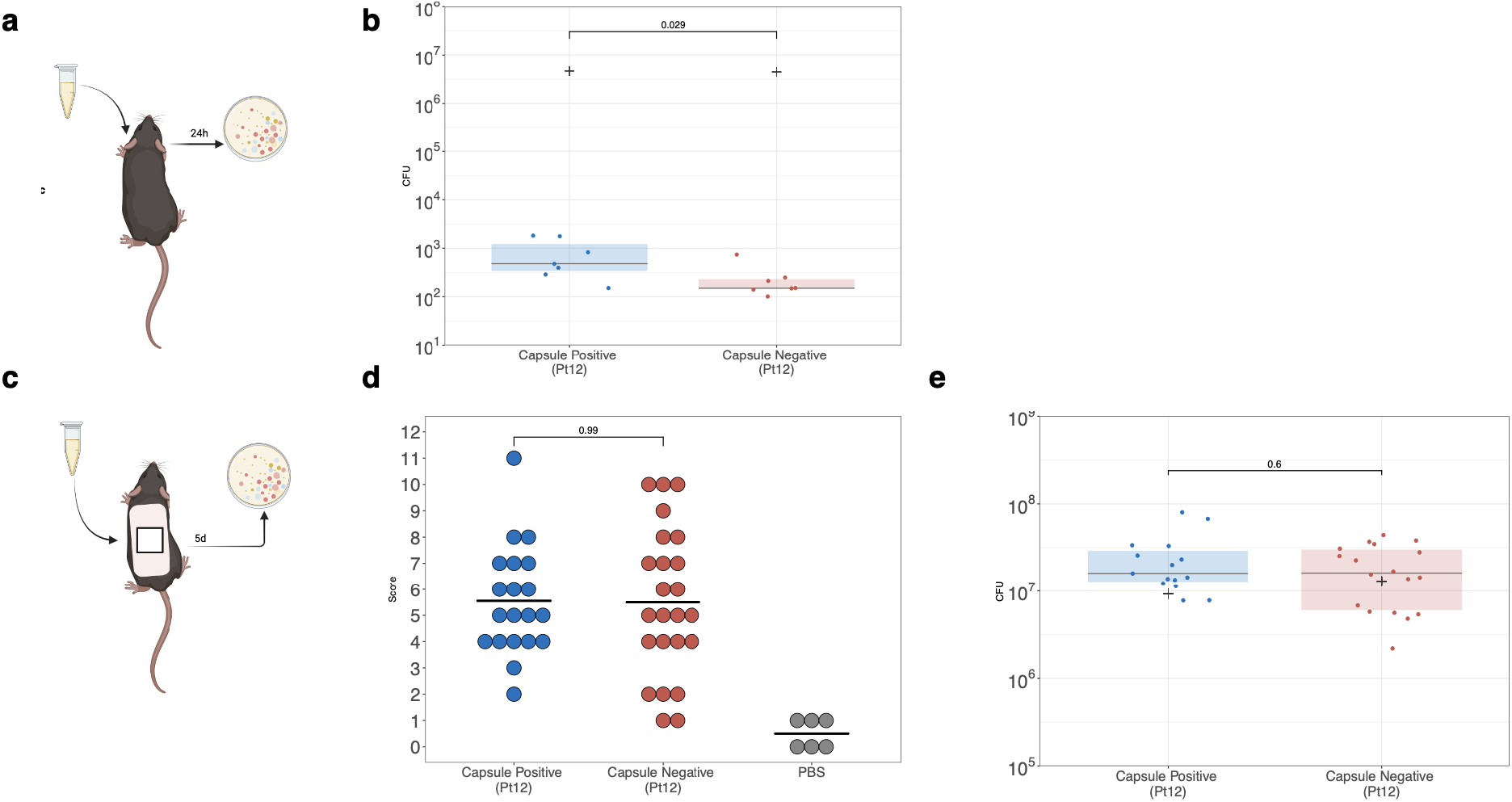
Capsule absence in clinical *S. aureus* strains does not enhance virulence or colonization in *in vivo* mouse models. **a.** Colonization / adherence was assessed using a simplified version of a previously published model ^1^. Bacterial strains were grown overnight, subcultured to log phase growth and washed to achieve normalized OD and applied to the ears. Bacterial load was quantified by CFU counting on ChromAgar after harvesting, homogenizing, and culturing tissue 24 h after colonization. **b.** The average recovered CFU from both ears is significantly lower (p=0.029, Wilcoxon Rank Sum) in mice treated with the capsule negative strain 059-AI7. **c.** Differences in virulence among capsular and acapsular strains was assessed using a previously published model of epicutaneous infection ^53^. **d.** The composite score represents the sum of individual assessments of erythema (0-3), oedema (0-3), thickness (0-3) and scale (0-3), which demonstrates no difference between capsule and acapsular strains. **e.** Bacterial load recovered from infected skin is equivalent among clinical isolate strains. Cross represents average inoculum across three replicates, n = 5-8 mice per group per replicate. All experiments are performed using capsular (048-RI3) and acapsular (059-AI7) isolates from patient 12 and compared using a Wilcoxon rank test.

**Supplementary Figure 2.**
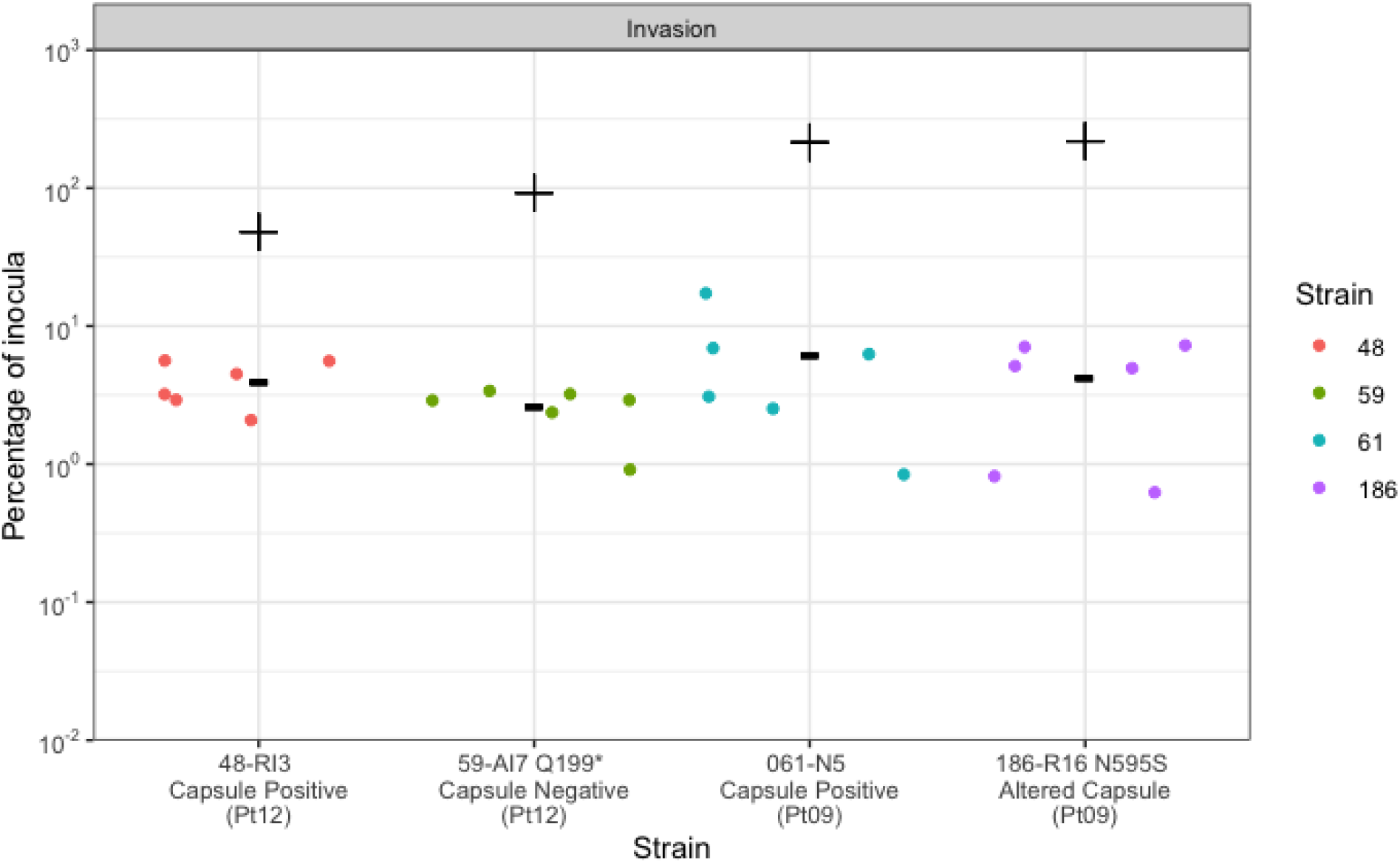
Capsule positive and negative strains invade human epithelial cells at comparable rates. The percentages of log phase and overnight inocula recovered from capsule positive and negative isolates from patient 12 and patient 09 were comparable between patient strains. Crosses represent MOI.

